# Symptomatic and Neurotrophic Effects of GABAA Receptor Positive Allosteric Modulation in a Mouse Model of Chronic Stress

**DOI:** 10.1101/2021.03.22.436517

**Authors:** Ashley Bernardo, Philip Lee, Michael Marcotte, Md Yeunus Mian, Sepideh Rezvanian, Dishary Sharmin, Aleksandra Kovačević, Miroslav Savić, James M. Cook, Etienne Sibille, Thomas D. Prevot

## Abstract

Chronic stress is a risk factor for Major depressive disorder (MDD), and in rodents, it recapitulates human behavioral, cellular and molecular changes. In MDD and after chronic stress, neuronal dysfunctions and deficits in GABAergic signaling are observed and responsible for symptom severity. GABA signals predominantly through GABAA receptors (GABAA-R) composed of various subunit types that relate to downstream outcomes. Activity at α2-GABAA-Rs contributes to anxiolytic properties, α5-GABAA-Rs to cognitive functions, and α1-GABAA-Rs to sedation. Therefore, a therapy aiming at increasing α2- and α5-GABAA-Rs activity, but devoid of α1-GABAA-R activity, has potential to address several symptomologies of depression while avoiding side effects. This study investigated the activity profiles and behavioral efficacy of two molecules enantiomers of each other (GL-II-73 and GL-I-54), separately and as a racemic mixture (GL-RM), and potential disease-modifying effects on neuronal morphology. Results confirm GL-I-54 and GL-II-73 exert positive allosteric modulation at the α2-, α3-, α5-GABAA-Rs and α5-containing GABAA-Rs, respectively, and have anti-depressant and pro-cognitive effects independently. Using unpredictable chronic mild stress (UCMS) in male and female mice (n=12/group), we show that acute and chronic administration of GL-RM combined the anti-depressant and pro-cognitive effects of each enantiomer, although at lower doses avoiding sedation. Morphology studies showed reversal of spine density loss caused by UCMS after chronic GL-RM treatment at apical and basal dendrites of the PFC and CA1. Together, these results support using a racemic mixture with combined α2-, α3-, α5-GABAA-R profile to reverse chronic stress-induced mood symptoms, cognitive deficits, and with anti-stress neurotrophic effects.

## Introduction

Chronic stress is a risk factor for psychiatric disorders including Major Depressive Disorder (MDD), displaying mood and cognitive symptoms [1–3]. Interventions to treat depression and other stress-related psychopathologies have focused on the monoaminergic system [4] with limited efficacy on mood symptoms, no efficacy on cognitive symptoms, and no effect on structural features of brain cells, highlighting unmet clinical needs. Recent studies have focused on the glutamatergic system [5], with promising effect on mood, cognition and cell structure [6, 7]. Similar efficacies were reported with compounds targeting the GABAergic system [8].

Mood and cognitive symptoms result from cellular dysfunctions, altered communication between cells and neuronal atrophy [9–13]. Transcranial magnetic stimulation in MDD patients shows reduced GABAergic function (i.e., cortical inhibition) [14, 15], contributing to impaired excitation/inhibition balance in MDD [16, 17]. Reduced GABA levels are reported in the occipital cortex [18–24], prefrontal cortex (PFC) [25] and anterior cingulate cortex [18, 25–27]. In the PFC, synaptic densities and expression of synaptic function-related genes are reduced [12, 28]. Chronic stress-related disorders [13, 29] and animal models report similar findings [30–32] and demonstrate critical links between prefrontal and hippocampal functions [33]. Chronic stress reduces dendritic length and spine density in the PFC and hippocampus (HPC), likely contributing to cognitive deficits. Of approved treatments, only the NMDA receptor antagonist, ketamine, has shown efficacy at increasing spine density and dendritic complexity in mice, along with antidepressant properties [34]. The neurotrophic effects of ketamine are hypothesized to act through BDNF-TrkB signaling while its fast-acting antidepressant properties are suggested to be initially mediated by activity on GABAergic neurons followed by long-term GABA and glutamate changes [35], highlighting a potential role for GABA in these therapeutic effects.

GABA signals through ionotropic GABAA receptors (GABAA-Rs) and metabotropic GABAB receptors. GABAA-Rs are pentameric ion channels, formed by a combination of 19 subunit subtypes [36]. Drugs acting on GABAA-Rs exist (benzodiazepines or imidazodiazepines), but their use is limited because of side effects (sedation) due to broad activity at several receptor subunits, including α1-GABAA-Rs [37–39]. Studies demonstrated α2-GABAA-Rs, strongly present in the amygdala [40–42] and HPC [40, 43, 44], are linked to regulation of anxious and depressive phenotypes [45–49]. The role of α3-GABAA-Rs are not fully characterized, but appear linked to behavioral despair in mice [50]. Other studies link α5-GABAA-Rs with cognitive performances [51–53], aligned with their preferential expression in cognitive processing regions (HPC and mPFC) [40, 54]. Our group and others showed targeting α5-GABAA-Rs with a positive allosteric modulator (PAM) reverses stress-[52] and age-[55, 56] related cognitive deficits, and chronically reverses age-related neuronal atrophy [53]. Potentiation of α2- and α5-GABAA-Rs, while limiting potentiation of α1-GABAA-R, is hypothesized to improve mood and cognitive symptoms while reducing side effects [37, 57], altogether suggesting potential for treatment of symptomatic and morphological alterations in chronic stress and MDD.

Previously, we showed the α5-PAM GL-II-73, has pro-cognitive, anxiolytic and anti-depressant properties at 10mg/kg IP but not at 5mg/kg IP [52], and neurotrophic effects at 30mg/kg PO [56] in mice. However, it remained unknown if its enantiomer, GL-I-54, has similar or complementary properties. Here, we investigated the similarities and/or complementarities of receptor selectivity and behavioral efficacy profile of GL-I-54 and GL-II-73 in stressed mice. We then investigated the symptomatic and neurotrophic potentials of a racemic mixture of both compounds in a mouse model of chronic stress.

## Materials and Methods

Details provided in the supplementary material.

### Compounds

GL-I-54 and GL-II-73 were synthesized as in [58], with >99% chemical and optical purity. GL-RM is a 1:1 mix of each compound.

### Electrophysiology

Electrophysiological recordings used HEK-293T cell line transfected with full-length cDNAs for human GABAA-R subtypes α(1/2/3/4/5)β3γ2 [59, 60]. PubMed Protein Sequence repository for GABAA-R subunits reports the binding pocket formed by α1/2/3/5 and γ2 is 100% conserved between mouse and human, predicting similar activity and selectivity across species. Current recordings were performed in absence or presence (5μM) of GABA. GL-I-54 or GL-II-73 was applied at 0.033-33.33μM concentrations.

### Animals

Six independent cohorts of 8-week old C57BL/6 mice (50% female) were obtained from Military Medical Academy (Belgrade, Serbia) or Jackson Laboratories (Stock#000664; MA, USA), measuring: pharmacokinetics (#1; n=25), individual enantiomers on elevated plus maze and forced swim test (#2; n=24), effect of GL-I-54 on Y-maze (#3; n=42), GL-I-54 side effects in the rotarod (#4, n=18), and acute (#5; n=36) and chronic (#6; n=36) efficacy of GL-RM. Animals were individually housed and maintained on a 12 hour light/dark cycle (7:00 ON, 19:00 OFF), with *ad libitum* food and water.

### Ethical Statement

All animal work was completed in accordance with Ontario Animals for Research Act (RSO 1990, Chapter A.22), Canadian Council on Animal Care (CCAC), Ethical Council for the Protection of Experimental Animals of the Ministry of Agriculture, Forestry and Water Management of the Republic of Serbia, and was approved by the Institutes’ Animal Care Committees.

### Drug Administration

GL-I-54 (1, 3, 5, or 10mg/kg), GL-II-73 (5mg/kg) or GL-RM (10mg/kg) were diluted in vehicle solution (85% distilled H2O, 14% propylene glycol (Sigma Aldrich) and 1% Tween 80 (Sigma Aldrich)) and administered intraperitoneal (IP), 30min before testing. For oral administration, GL-RM was prepared in tap water (30mg/kg) and given through drinking water, considering 6mL average daily fluid intake/animal.

### GL-I-54 Pharmacokinetics

Mice were treated IP with 3mg/kg GL-I-54 and euthanized at different time points (5-720 min post-injection) for brain and plasma quantification of ligand by ultraperformance LC-MS/MS (cassette dosing) [61]. In vitro hydrolytic plasma stability of GL-I-54 was tested in vitro at 37°C, utilizing blank mouse plasma spiked with GL-I-54 and internal standard, as in [62].

### Plasma Protein and Brain Tissue Binding Studies

A rapid equilibrium dialysis assay determined the free fraction of GL-I-54 in mouse plasma and brain tissue as in [63]. GL-I-54-free brain concentrations were calculated by multiplying the total brain concentrations with the appropriate free fractions determined by rapid equilibrium dialysis.

### Liver Microsomal Assay

Metabolic stability was tested in C57BL/6 Mouse liver microsomes at 2μM and 0.5mg/mL of matrix concentration, as in [64]. NADPH solution, Microsomes/S9 and compounds were added to assay plates. Plates were quenched at T0, 30, 60 and 120 min and assessed in refrigerated LC-MS/MS autosampler. Intrinsic hepatic clearance calculations used 45mg microsomes/g of liver and 87.5g liver/kg of body weight for mouse.

### Stress Paradigms

Chronic restraint stress (CRS) was used with cohorts #2 and #3, consisting of placing mice in a 50mL Falcon™ Tube for 1hr twice daily for 1 week. Cohorts #5 and #6 were subjected to Unpredictable Chronic Mild Stress (UCMS), using randomized mild stressors (twice daily) over 6 weeks. Weeks with behavioral testing applied one stressor per day after the behavioral testing (2 weeks). UCMS animals were housed in a separate room without environmental enrichment to exacerbate effects of other stressors.

### Behavioral Tests

Side effects were measured in the rotarod test in cohort #4, assessing locomotor coordination on an accelerating rod. Latency to fall was measured over 6 trials. Mice in cohorts #2 were tested in the elevated plus maze (EPM), measuring exploration of open arms for 10 min as a proxy of anxiety [65], and in the forced swim test (FST), measuring time spent immobile for 6 min as a proxy of antidepressant-like effect. In cohort #3, mice were exposed to CRS to induce a working memory in the Y maze test, and the capacity of reversal of GL-I-54. In cohorts #5 and #6, mice were subjected to UCMS, and each animal’s coat state was scored (0=well-groomed, smooth coat, 1=soiled coat or bald patches) weekly for seven coat regions, as in [66]. Behavioral screen included: Anxiety-like behavior (EPM, open-field, OF; Novelty Suppressed Feeding, NSF; and the Phenotyper Test [65], PT), anhedonia (Sucrose Consumption; SC), antidepressant-like activity (FST) and cognitive function (Y-maze alternation task [52, 56, 67]).

### Tissue Collection and Golgi Staining

Cohort #6 was euthanized by cervical dislocation 24hr after last behavioral test. Brains were immersed in Golgi-Cox staining solution, and assigned a unique identifier (4 brains/group). Brains were shipped to NeuroDigiTech (San Francisco CA, USA) for sectioning (100μm thickness), mounting and blind quantification of basal and apical dendrites of PFC and CA1 pyramidal cells using NeuroLucida v10 software. Neurons (n=6/animal) were analyzed for dendritic length, spine number and spine density [68].

### Statistics

Statistical analyses used GraphPad Prism 9. Electrophysiology data and compounds activity at GABAA-R subunits were compared to 100% using t-tests. Other analyzes used two-way ANOVA with concentration and subunit as co-factors, and Bonferroni post-hoc tests. Behavioral analyzes used one-way or two-way ANOVA, and repeated measure ANOVA as relevant. Fishers PLSD tests were used for post-hoc analyses. Spearman ranks correlation was used between morphological features and behavioral outcomes. Statistical significance was considered p<0.05. Data are presented as mean ± SEM.

Z-scores were calculated to assess consistency of anxiety-like and depressive-like behavioral phenotypes across tests, referred to as z-emotionality using averaged z-scores of behavioral tests as in [69]. Full calculations in supplementary.

## Results

Statistical significances are provided in **Supplementary Tables**.

### GL-I-54 and GL-II-73 exhibit different GABAA-R potentiation profiles but similar metabolic stability

Both enantiomers were confirmed PAMs, not agonists (**Supplementary Fig. 1 and Fig. 1A-B**), acting only in the presence of GABA. GL-I-54 significantly increased potentiation at α2-GABAA-Rs (>0.03μM), α3-GABAA-Rs (>0.33μM), α1-GABAA-Rs (>1μM) and α5-GABAA-Rs (0.03, 0.33 and 3.33μM; **Fig. 1A**), with preferential activity at α3- and α2-over α1-, α4- and α5-GABAA-Rs **(Supplementary Table 1A**).

**Figure 1.**
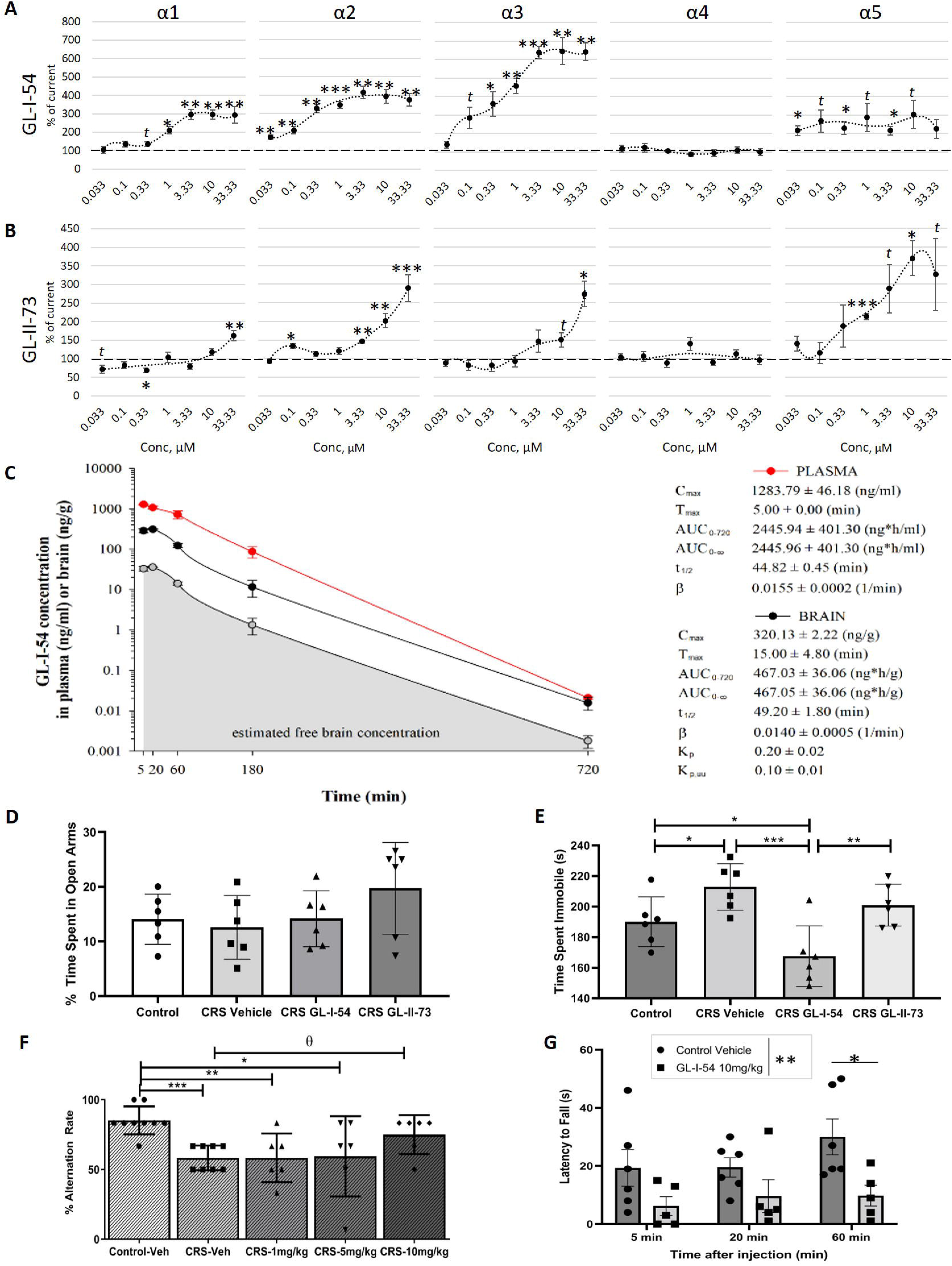
Electrophysiological, pharmacokinetic and behavioral profiles of GL-II-73 and GL-I-54. Electrophysiological recordings were obtained from HEK-293T cells transfected with full-length cDNA for human GABAA receptor subtypes α1β3γ2, α2β3γ2, α3β3γ2, α4β3γ2 or α5β3γ2, in presence of GL-I-54 (**A**) or GL-II-73 (**B**), and in the presence of GABA in the medium (5μM). Pharmacokinetic profile of GL-I-54 was also examined (**C**). Plasma, brain and free brain concentration-time profile of GL-I-54 after intraperitoneal cassette administration of 3 mg/kg dose in male C57BL/6 mice (n = 3 per time point). C_max_, maximum concentration in plasma or brain; T_max_, time of maximum concentration in plasma or brain; AUC_0–720_, area under the plasma or brain concentration-time curve from 0 to 720 min; AUC_0-∞_, area under the plasma or brain concentration-time curve from 0 to extrapolated infinite time; t_1/2_, elimination half-life from plasma or brain; β, elimination constant rate from plasma or brain; K_p_, brain-to-plasma partition coefficient (K_p_ = AUC_0-∞_, brain/AUC_0-∞_, plasma); K_p,uu_, ratio of unbound brain to unbound plasma drug concentrations (K_p,uu_ = K_p_ × unbound fraction in brain/unbound fraction in plasma). GL-I-54 and GL-II-73 were then tested in the elevated plus maze and forced swim test at the dose of 5mg/kg, in mice previously exposed to chronic restraint stress (CRS). Drugs were administered IP, 30min prior to testing. Percent time spent in the open arms (**D**) showed no difference between groups. Time immobile (**E**) showed more time spent immobile in the CRS-Vehicle group, and less time immobile in animals receiving GL-I-54, compared to CRS-Vehicle. GL-II-73 did not show an effect. GL-I-54 was tested in the Y-maze task, assessing working memory (**F**). Animals subjected to CRS and receiving vehicle showed a significant decrease in alternation rate, suggesting a working memory deficit. Animal subjected to CRS and receiving the highest dose of GL-I-54 showed significant increase in alternation rate, suggesting reversal of working memory deficits induced by CRS. Finally, independent mice were tested in the rotarod (**G**; N=5 Control Vehicle and N=6 GL-I-54). Mice were trained to maintain themselves on a rotating rod (rotarod) for 3 trials. Then, they were injected with GL-I-54 at 10 mg/kg, and tested 5 min, 20 min and 60 min past injection time. Latency to fall from the rod was recorded, and showed significant reductions in latency to fall in mice receiving GL-I-54. All values are represented as mean ± standard error of the mean. *p<0.05, **p<0.01, ***p<0.001 compared to 100 % in A-B) or to Control, Control/Vehicle in D-G. θp<0.05 compared to CRS-Vehicle.

GL-II-73 significantly increased potentiation at α5-GABAA-Rs (>1μM), α2-GABAA-Rs (3.33μM), α3-GABAA-Rs (33.33μM) and α1-GABAA-Rs (33.33μM; **Fig. 1B**), with preferential activity at α5-GABAA-Rs over α1-GABAA-Rs (>0.3μM), α2-GABAA-Rs (between 3.3 and 10μM), α3-GABAA-Rs (between 1 and 10μM) and α4-GABAA-Rs (>0.33μM) (**Supplementary Table 1B**), confirming data previously published [52].

### Pharmacokinetic profile, brain penetrance and stability

Pharmacokinetic profile of GL-I-54 dosed at 3mg/kg in mice is represented in **Fig. 1C**. Accounting for the doses administered in respective studies, GL-I-54 and GL-II-73 exhibit comparable pharmacokinetic patterns. As a representative example, 20min after the 3mg/kg dose, GL-I-54 achieved 311.75±6.72ng/g, while GL-II-73 reached 264.61±38.22ng/g (**Supplementary Fig. 2)**. After appropriate adjustments (for the 5mg/kg dose, free fraction, brain tissue density and unit), these close-to-maximum concentrations achievable in the mouse brain correspond to the 0.100-0.330μM concentration range in electrophysiological recordings (calculation in **Supplementary Fig. 2**). This implies that, in acute behavioral experiments with 5mg/kg dose, GL-I-54 likely potentiates α_1_, α_2_, α_3_ and α_5_ GABAA-Rs, however, α_1_ potentiation is mild. 5mg/kg of GL-II-73, on the other hand, is fairly silent on all GABAA-Rs, with the potential to mildly activate α_2_ and α_5_ receptors; on α_1_ and α_3_ receptors, nevertheless, it behaves as a null modulator. Plasma and brain free fractions of GL-I-54 were 22.57% and 11.49%, respectively, while, as previously reported for GL-II-73 were 20.39% and 12.14% [52], respectively. Brain to plasma ratio (K_p_) evaluates brain penetrance (penetrant when K_p_>0.04). GL-I-54 is brain penetrant (K_p_=0.20), similarly to GL-II-73 (K_p_=0.30), which is further supported by the ratio of unbound brain to unbound plasma ligand concentrations values, K_p,uu_ (0.10 vs 0.18, respectively). GL-I-54 displayed high in vitro metabolic stability, although lower compared to GL-II-73; after 4hr of incubation in mouse plasma, the fraction of remaining intact ligand was 78.41% (vs 98.88% of GL-II-73 [52]). In the liver microsomal assay, GL-I-54 showed a 198min half-life with 70% remaining after 2hr. Intrinsic hepatic clearance was 27.6mL/min/kg. GL-II-73 showed a longer half-life, 256min, with 75.6% remaining after 2hr. Intrinsic hepatic clearance was 21.3mL/min/kg.

### Acute administration of GL-I-54 displays antidepressant and procognitive effects in the chronic retrain stress model with locomotor side effects at higher dose

In mice subjected to CRS, anxiolytic and antidepressant properties of GL-I-54 (5mg/kg) and GL-II-73 (5mg/kg) were tested using the elevated plus maze (EPM), forced swim test (FST). Pro-cognitive properties were tested in the Y-maze (5mg/kg and 10mg/kg). After CRS, GL-I-54 and GL-II-73 had no effect on percent time in open arms of the EPM (**Fig. 1D**). In the FST, CRS increased time spent immobile, which was significantly reversed by GL-I-54 (**Fig. 1E**), suggesting antidepressant potential. In the Y-maze, CRS induced an alternation rate deficit that was reversed by GL-I-54 at 10mg/kg but not 5mg/kg (**Fig. 1F**), suggesting the capacity of reversing memory deficits. Locomotor side effects of GL-I-54 at 10mg/kg were tested in the rotarod (**Fig. 1G**). Decreasing latency to fall 60min after injection was found, suggesting sedation at this dose.

### Acute administration of GL-RM displays antidepressant and procognitive effects in the UCMS model

Based on the selective effect on mood and no effect on working memory at 5mg/kg for GL-I-54 or GL-II-73 alone, we tested potential additive and low side effects using a 1:1 racemic mixture of 5mg/kg each, called GL-RM, in the UCMS model (**Fig. 2A**), with a final GL-RM dose of 10mg/kg. Weekly monitoring showed UCMS deteriorated coat state, not reversed by acute treatment, and weight gain remained unaffected (**Supplementary Fig. 3**).

**Figure 2.**
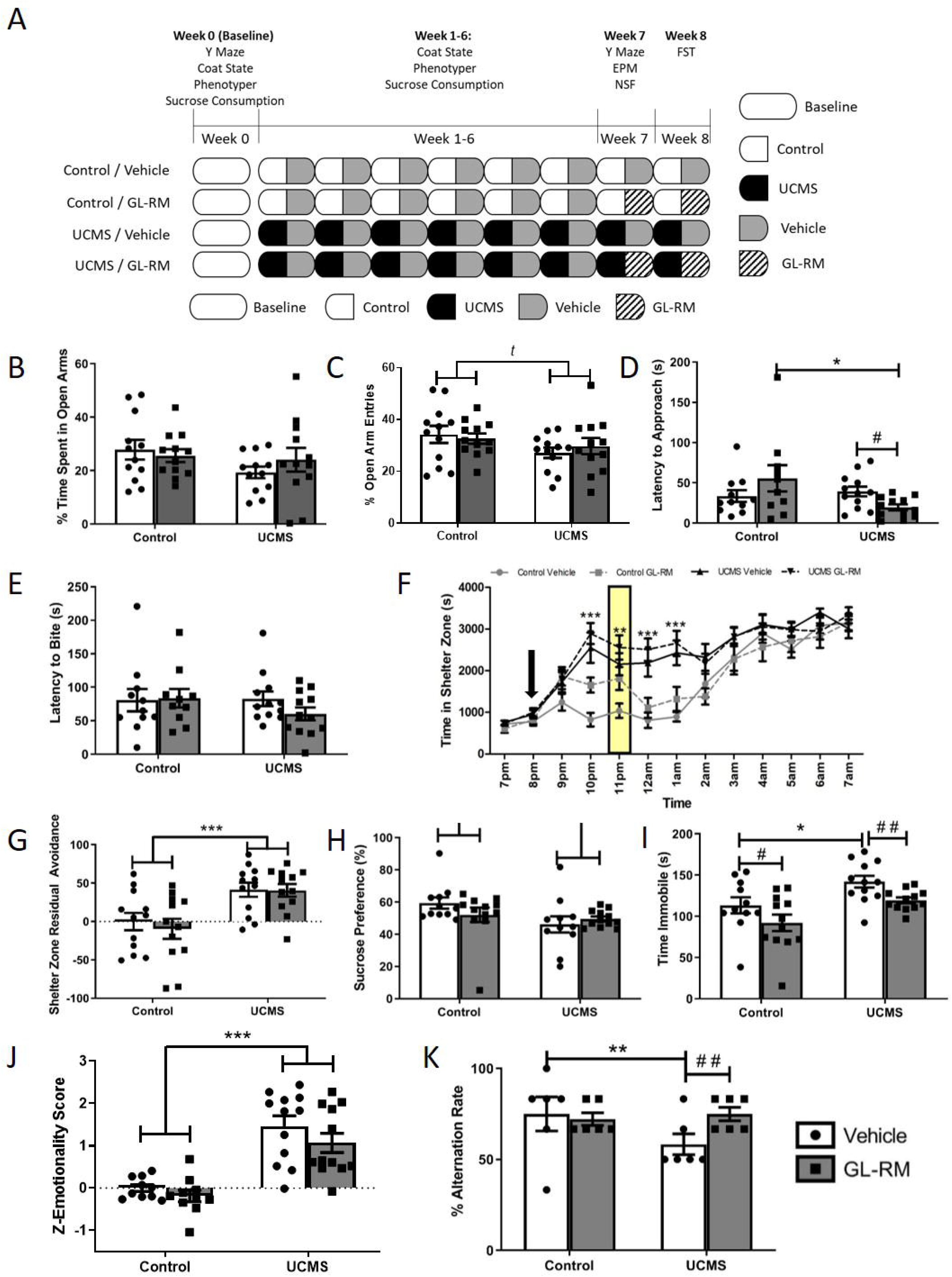
Effect of acute treatment of GL-RM on anxiety, emotionality and working memory deficits in mice subjected to chronic stress. Male and female mice were tested at baseline in the Y-maze, the phenotyper and the sucrose consumption test prior to being subjected to 6 weeks of UCMS (**A**). Weekly, mice were tested in the phenotyper test, the sucrose consumption and their weight and coat state were measured. After 6 weeks of UCMS, acute injections were performed 30 minutes prior to behavioral testing. In the elevated plus maze, time spent (**B**) and entries (**C**) in the open arms were measured, with no significant effect of UCMS, treatment or interaction. In the novelty suppressed feeding test, latency to approach (**D**) and latency to bite (**E**) were assessed. Statistical analyses showed that acute GL-RM treatment reduces latency to approach in mice subjected to UCMS. In the phenotyper test (**F**), mice were placed in the box overnight, where a stressful stimulus was applied at 11pm, for 1hr. Time spent in the shelter zone showed that mice subjected to UCMS spent more time in the shelter than control mice. Calculating a residual avoidance score (**G**), statistical analyses showed a significant increase in residual avoidance in mice subjected to UCMS. In the sucrose consumption test (**H**), mice subjected to UCMS showed a significant decrease in preference to sucrose. In the forced swim test (**I**), mice subjected to UCMS showed increased immobility, while mice treated with GL-RM showed a reduction in immobility. Combining the individual score into a global z-score (**J**), statistical analyses confirmed a significant impact of UCMS, with reduced effect of GL-RM. Finally, mice were tested in the Y-maze (**K**), where statistical analyses showed altered alternation with UCMS, which is reversed by acute GL-RM treatment. ^*t*^p<0.1, *p<0.05, **p<0.01, ***p<0.001 effect of UCMS, #p<0.05, ##p<0.01 effect of GL-RM

In the EPM, there was no effect of UCMS or treatment on percent time in open arms (**Fig.2B**). There was a trend level effect of stress reducing the percent of entries into open arms but no effect of treatment (**Fig.2C**). NSF latency to approach found a significant interaction between UCMS and treatment, due to reduced latency in UCMS-GL-RM treated animals compared to Control-GL-RM and compared to UCMS-vehicle (**Fig.2D**), suggesting anxiolytic properties. Latency to bite was unaffected by UCMS or treatment (**Fig. 2E**). Phenotyper test and associated measure of residual avoidance used in Prevot et al. [65] assess anxiety-like behavior weekly. The residual avoidance score captures the time spent in the shelter, after a 1hr-light challenge, measuring residual avoidance of the previously-lit zone. Prior to UCMS, there was no group difference (**Supplementary Fig. 4A-B**). Upon UCMS, residual time spent in the shelter following an acute light challenge was increased (**Fig. 2F-G**; **Supplementary Fig. 4C-G**). On week 6, acute GL-RM treatment did not affect time in the shelter after the light challenge (**Fig. 2F**). Sex-dependent analyses showed males spent more time in the shelter than females (data not shown).

Anhedonia-like and antidepressant-like behaviors were evaluated using the SC and FST tests, respectively. At week 6, sucrose preference was significantly reduced by UCMS but unaffected by GL-RM (**Fig. 2H**), suggesting a lack of effect on anhedonia. In the FST, UCMS increased time immobile, which was reduced by GL-RM (**Fig. 2l**), suggesting antidepressant properties. Z-scoring of behavioral emotionality confirmed the effect of UCMS, and a limited efficacy of acute GL-RM treatment (**Fig. 2J**). In contrast, the Y-maze spatial working memory deficit induced by UCMS was significantly reversed by acute GL-RM treatment (**Fig. 2K**).

Altogether, these results showed that acute administration of GL-RM has antidepressant and procognitive potential in the UCMS model, while anxiety measures are inconclusive.

### Chronic GL-RM conserves antidepressant and pro-cognitive properties in the UCMS model

The effects of chronic GL-RM administration were investigated in mice subjected to UCMS (**Fig. 3A**). Altered coat state and reduced weight gain with UCMS was significant from week 1-6, with no effect on chronic GL-RM (**Supplementary Fig. 5**).

**Figure 3.**
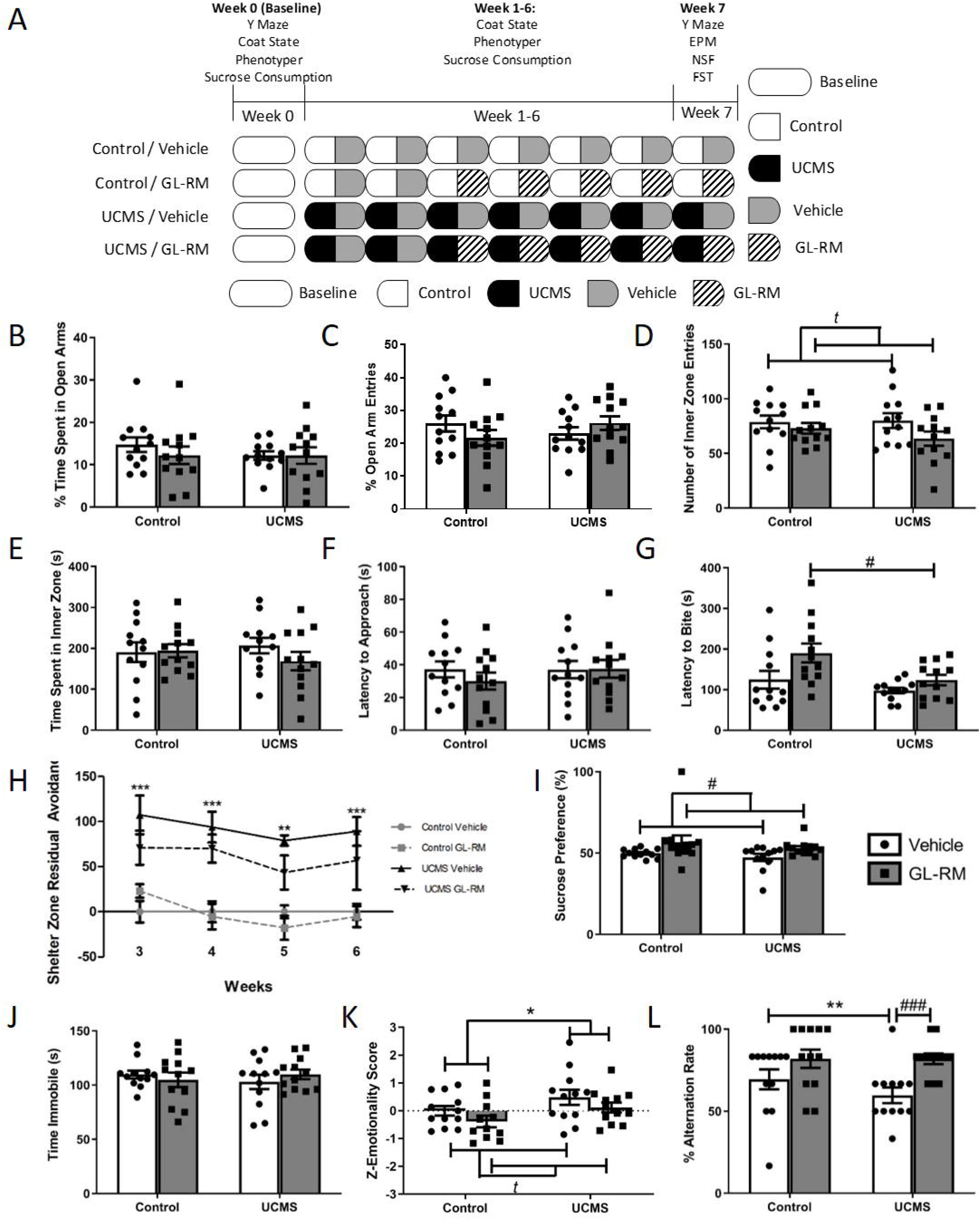
Effect of chronic treatment of GL-RM on anxiety, emotionality and working memory deficits in mice subjected to chronic stress. Male and female mice were tested at baseline in the Y-maze, the phenotyper and the sucrose consumption test prior to being subjected to 6 weeks of UCMS (**A**). After 3 weeks of UCMS, chronic treatment with GL-RM in the drinking water was initiated, for a total of 4 weeks. Weekly, mice were tested in the phenotyper test, the sucrose consumption and their weight and coat state were measured. After 6 weeks of UCMS, and 3 weeks of treatment, mice were tested in the elevated plus maze. Time spent (**B**) and entries (**C**) in the open arms were measured, but did not show statistical differences. Mice were also tested in the open field for the time spent (**D**) and number of entries (**E**) in the inner zone. Again, statistical analyses did not reveal any effect of UCMS nor treatment. In the novelty suppressed feeding test, latency to approach (**F**) and latency to bite (**G**) were assessed. Mice were tested in the Phenotyper weekly, and the residual avoidance scores from week 3 to 6 were analyzed, since the treatment was onboard during these testing periods (**H**). In the sucrose consumption test (**I**), mice receiving chronic GL-RM showed a significant increase in preference to sucrose. In the forced swim test (**J**), statistical analyses did not reveal significant differences between groups. Combining the individual score into a global z-score (**K**), statistical analyses confirmed a significant impact of UCMS at increasing emotionality, with a trend level effect ofchronic GL-RM reducing emotionality. Finally, mice were tested in the Y-maze (**L**), where statistical analyses showed altered alternation with UCMS, which is reversed by acute GL-RM treatment. *p<0.05, **p<0.01, ***p<0.001 effect of UCMS, ^*t*^p<0.1, #p<0.05, ###p<0.001 effect of GL-RM

There was no effect of UCMS or treatment in the EPM, OF and NSF latency to approach (**Fig. 3B-F**). Latency to bite was decreased in UCMS-GL-RM mice compared to Control-GL-RM mice (**Fig. 3G**). In the Phenotyper test (**Fig. 3H** and **Supplementary Fig. 6**), UCMS increased residual avoidance in the shelter zone, while chronic GL-RM was trending to decrease it.

Sucrose preference was unaffected by UCMS, but chronic GL-RM increased preference (**Fig. 3I**), suggesting improvement of anhedonia-like behavior. In the FST, the time spent immobile was unaffected by UCMS or treatment (**Fig. 3J**). UCMS significantly increased z-emotionality, with a trend level effect of chronic GL-RM reducing z-emotionality (**Fig. 3K**).

In the Y-maze, UCMS decreased percent alternation rate (Control-Vehicle vs UCMS-Vehicle) and chronic GL-RM significantly increased percent alternation rate between UCMS-vehicle and UCMS-GL-RM groups (**Fig. 3L**), confirming efficacy at reversing stress-induced memory deficits.

Results suggest that chronic GL-RM may contribute to reducing z-emotionality, with discrete effects on individual mood symptoms, and confirm its effect at reversing UCMS-induced working memory deficits.

### Chronic GL-RM administration prevents neuronal morphology deficits induced by UCMS

We quantified morphological changes in apical and basal dendrites of PFC pyramidal neurons (**Fig. 4A-C**, and **Supplementary Fig.7**). Dendritic length was not affected by UCMS or treatment, but spine count and spine density were significantly lower in UCMS compared to control mice (**Fig. 4D-E**, and **Supplementary Fig.7**). Such decreases were reversed by chronic GL-RM in basal and apical segments, with spine density in apical PFC dendrites of GL-RM treated UCMS animals being significantly higher than UCMS animals and not significantly different from controls (**Fig. 4D**).

**Figure 4.**
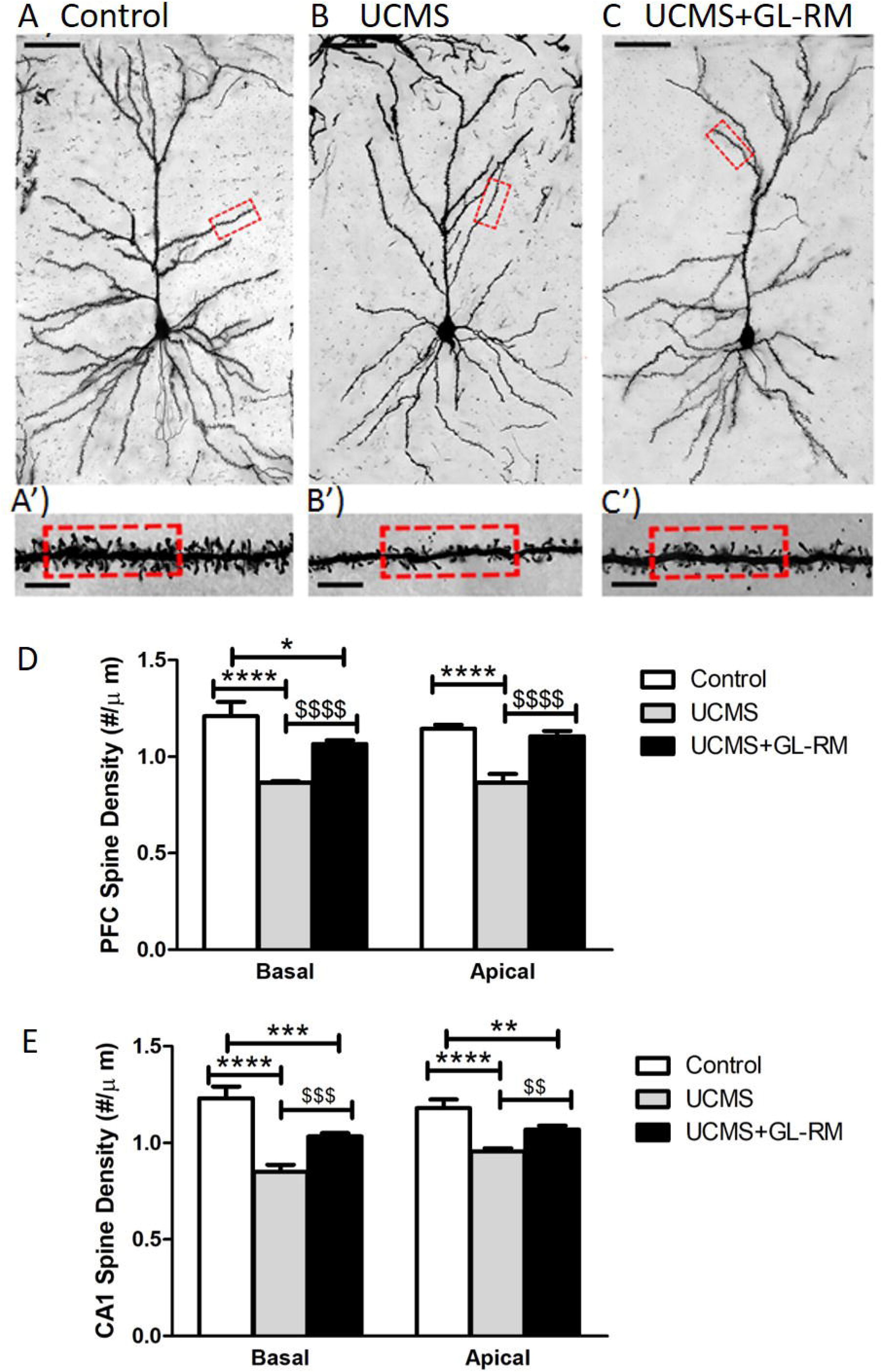
Chronic treatment with GL-RM reverses chronic-stress induced spine density reduction in the PFC and the CA1. After completion of the behavioral screening, mice were euthanized and brains were stained with Golgi-Cox solution. Pyramidal neurons (N=6 per mouse) from 4 mice per group (**A-C**) were analyzed for dendritic length, spine counts and spine density. Basal and apical spine densities were measured in the PFC (**D**) and the CA1 of the hippocampus (**E**). ANOVA in the basal and apical segments revealed significant differences between groups, in both brain regions. This difference was explained by a decrease in spine density in mice subjected to UCMS compared to Control mice that was partially reversed by chronic treatment with GL-RM. *p<0.05, **p<0.01, ***p<0.001, ****p<0.0001 compared to “Control”; $$p<0.01, $$$p<0.001, $$$$p<0.0001 compared to “UCMS”. Scale bar in (A-C) represents 50μm.

Similarly, significant UCMS effects on spine density and reversal by chronic GL-RM treatment were observed in CA1 pyramidal neurons (**Fig. 4E**, and **Supplementary Fig. 8**).

### Spine architecture in the PFC and CA1 are related to spatial working memory and emotionality

Spine densities in the PFC and CA1 were correlated to spatial working memory performance in the Y-maze (**Fig. 5 and Supplementary Table 5)**. In both regions, positive correlations were found at apical and basal segments, with lower alternation rates associated with lower spine densities in the PFC (**Fig. 5A–5B)** and the CA1 (**Fig. 5C-D**). No correlations were found for dendritic length or spine count in either region with Y-maze performance.

**Figure 5.**
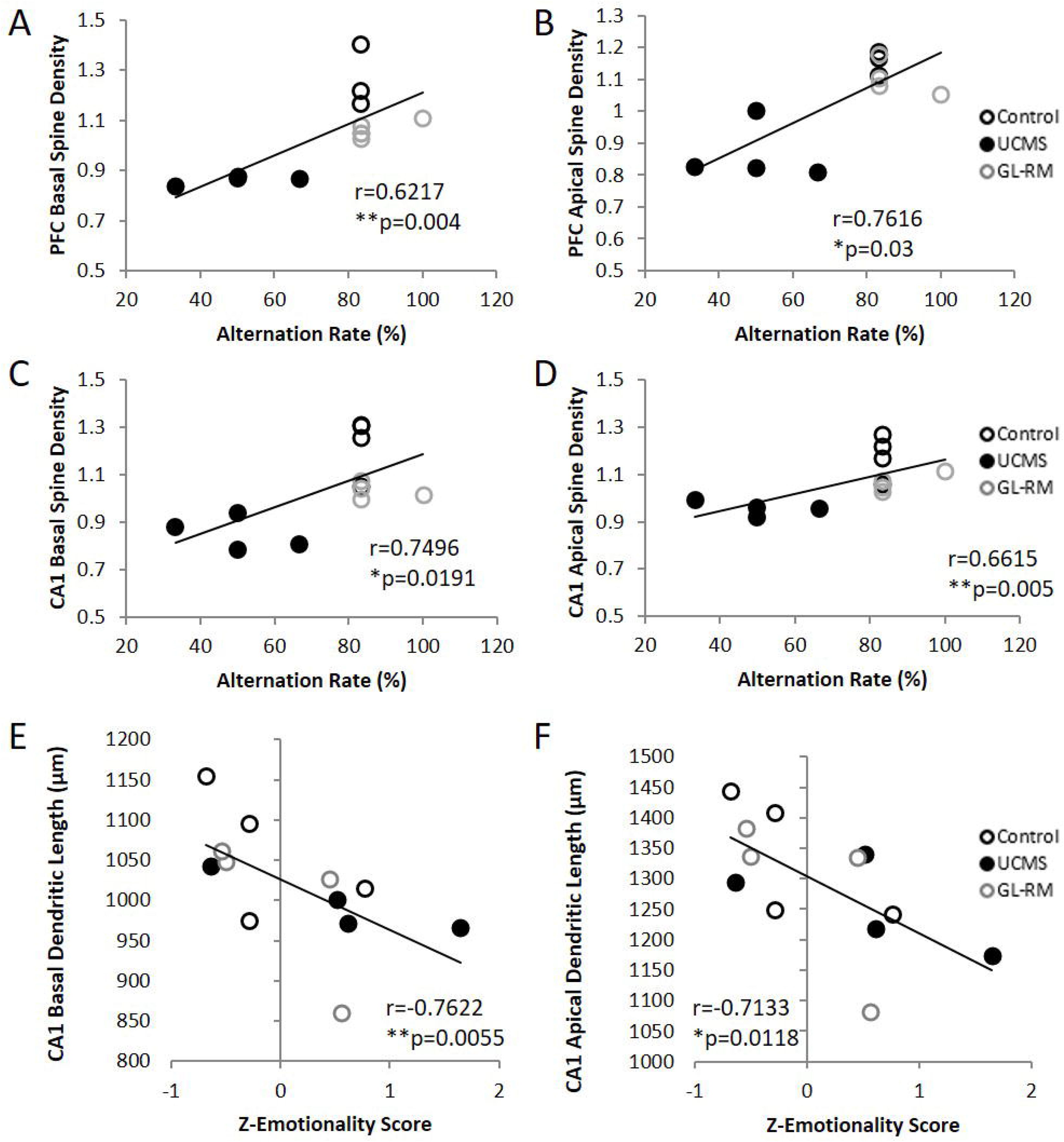
Morphological changes in the PFC and CA1 correlates with behavioral outcomes. Morphological features from the PFC and the CA1 were correlated to behavioral outcomes using the alternation rate (**A-D**) and the z-emotionality (**E-F**). Basal (**A**) and apical (**B**) spine density in the PFC correlated positively with the alternation rate, where higher density was associated with higher alternation rate, i.e. higher memory performances. Similar findings were reported when looking at basal and apical spine density from the CA1 (**C-D**). Basal and Apical dendritic length in the CA1 also correlated with z-emotionality, this time negatively (**E-F**), where higher dendritic length was associated with lower z-emotionality, i.e. lower anxiety- and depressive-like behaviors.

Correlations between morphological outcome and z-emotionality were calculated (**Supplementary Table 5)**. There were no significant correlations between z-emotionality and any morphological parameters in the PFC. In the CA1, increased dendritic length correlated with lower z-emotionality scores in basal and apical dendrites (**Fig. 5E-F**), with no correlations with spine count or spine density.

## Discussion

This study is based on observations that patients with psychopathologies related to chronic stress exposure experience mood and cognitive deficits, while current treatments display moderate-to-no efficacy for mood and cognition [1–3, 9]. We investigated the efficacy of two enantiomers with activity at α3-, α2-, α5-GABAA-Rs (GL-I-54) and α5-GABAA-Rs (GL-II-73) separately, and in combination (GL-RM) at alleviating such symptoms and neuronal atrophy in CRS and UCMS models. GL-I-54 showed antidepressant-like effects, no anxiolytic nor cognitive effects at low dose (5mg/kg), and reverses working memory deficits at a higher dose (10mg/kg), while causing slight sedation. GL-II-73 previously showed no effect on working memory at doses lower than 10mg/kg [52]. Acute and chronic treatment of a racemix combination of both enantiomers (GL-RM) improved stress-induced cognitive deficits. Acute GL-RM showed anti-depressant potential but no anxiolytic effects. Finally, chronic GL-RM treatment reversed the UCMS-induced reduction in spine densities in the PFC and CA1 of UCMS mice, with morphological measures correlating with improvement of emotionality and working memory.

### Racemic mixtures have therapeutic relevance by harnessing different enantiomer GABAA-R specificities

Therapeutic development often favors pure substances over racemic mixtures due to superior selectivity and reduced off-target activities [70, 71]. Racemic development becomes justified if therapeutic benefits of each enantiomer are similar, complimentary or devoid of side-effects and toxicity [72]. GL-RM harnessed the similar, yet unique selectivity profiles of GL-I-54 and GL-II-73 to target the GABAA-Rs known to be involved in anxiety, depression and cognition. GL-I-54 displays activity at α3-, α2-, α5-, α1-GABAA-Rs while GL-II-73 confirmed preferential activity at α5-GABAA-Rs [52], likely reflecting different interactions of each compound’s conformation in the binding pocket.

Racemic mixtures are not uncommon in the treatment of MDD [73, 74]. Fluoxetine [73, 75], citalopram [76] or ketamine [34] are clinically used racemic antidepressants [73, 77, 78]. Racemic R,S-ketamine has more efficacy than non-racemic S-Ketamine [79]. Racemic therapies can harness broader selectivity profiles to address multiple pathophysiological mechanisms but have the added risk of off-target side effects. *In vivo*, considering estimated free concentrations in the brain after a 5mg/kg dose, we found that concentration is slightly above 100nM. At this concentration, GL-I-54 may act as a PAM at non-α1 GABAARs, while GL-II-73 would mainly have only slight PAM activity at α2- and α5-GABAA-Rs, and be a null modulator at α1-GABAA-Rs. When combined together, low concentrations of GL-II-73 may be effective as an “on-site softener” of activity of GL-I-54, especially at α1-GABAA-Rs, limiting binding of GL-I-54 in the binding pocket and therefore limiting negative effects observed when potentiating this subunit, further suggesting GL-RM as an intervention for treatment of complete depression symptomatology.

### Targeting α2-, α3-, α5-GABRA-Rs with GL-RM displays superior anti-depressant and pro-cognitive effects at lower concentrations than individual enantiomers

Reconciling potentiation profiles with behavioral results, we compared GL-I-54 and GL-II-73 alone at 5mg/kg or 10mg/kg to GL-RM 10mg/kg (each enantiomer at 5mg/kg). At 5mg/kg, GL-I-54 has an antidepressant effect by reducing immobile time in the FST. Previously reported, GL-II-73 only reduced immobility time at 10 not 5mg/kg [52]. GL-RM reduced immobility time in the FST suggesting GL-RM retains antidepressant effects likely due to GL-I-54. Neither enantiomer shows clear anxiolytic effects alone or in GL-RM.

Interestingly, GL-RM combined the efficacy of both enantiomers for pro-cognitive effects. While neither enantiomer elicited pro-cognitive effects at 5mg/kg alone, their combined activity in GL-RM improved working memory deficits. This could be the results of an additive effect of the two compounds at GABAA-Rs, or a reduction of a putative amnestic effects caused by α1-GABAA-R null potentiation [80] with the use of a low GL-II-73 dose. Nevertheless, this finding demonstrates a clinical advantage for using GL-RM over either pure enantiomer at low concentrations, avoiding side effects and benefiting from combined profiles.

### Chronic GL-RM administration has neurotrophic effects

As in MDD, chronic stress leads to reduced synchronicity between the PFC and HPC, and dysfunctional information processing within cortical and hippocampal structures [81]. Dendritic length and spine density are reduced in the PFC and HPC of MDD patients and stressed mice [82], contributing to cognitive deficits. Our findings are consistent with other UCMS studies reporting neuronal atrophy and working memory deficits [83–85]. We previously showed that chronic administration of GL-II-73 alone remedies age-related spine loss within apical segments of pyramidal neurons, through α5-GABAA-R modulation confined to apical dendrites [56], because α5-GABAA-Rs are primarily located at apical dendrites [86–88] and GL-II-73 is preferentially selective for α5-GABAA-Rs. With GL-RM, neurotrophic effects were extended to basal segments in both PFC and HPC, suggesting a potential role of α2- and/or α3-GABAA-Rs in this neurotrophic effect. Previously, α5- and α2-GABAA-Rs have been implicated in dendritic outgrowth, spine maturation and synapse formation [89–91], thus their modulation by GL-RM may be responsible for spine density restoration at apical and basal dendrites.

While showing spine count improvements, it remains unclear whether spines are prevented from shrinking, or if they are generated *de novo*. Ketamine showed ability to overcome stress hormone-induced spine loss by a combination of restored spines and *de novo* spines, with some preference for *de novo* [92]. A similar combination may also be the case for GL-RM. Evidence that GL-RM has an effect at a pathophysiological level (not only symptomatic levels) is substantially valuable for translational efficacy within humans.

### Limitations

We tested GL-RM on a single cognitive domain, exploring others could add more insight. Regarding neurotrophic effects, we investigated the PFC and HPC for their regulation of stress-related and depressive symptoms. It could be valuable to investigate how GL-RM modulates connectivity in subnuclei of the amygdala, potentially decreasing the hyperactivity/increased connectivity reported after chronic stress [93, 94]. We used chronic stress models because they recapitulate behavioral and cellular changes observed in human depression [95–98]. While using the most commonly reported stressors [99], variability in anxiety-related outcomes limited clear conclusions on procedure and treatment, not common in behavioral studies [65, 100]. The Phenotyper test measuring context anxiety [65] provided insight to anxiolytic properties of GL-RM with better consistency. The discrepancy between EPM/NSF and Phenotyper results highlight potential biases from hands-on behavioral assays versus automatized approaches [101].

To conclude, the therapeutic relevance of a targeted PAM approach at GABAA-Rs is reinforced and the role of α5-GABAA-Rs in cognition is supported. At low-to-moderate dose, both compounds, used together, show promising effects for the treatment of mood and cognitive symptoms, as well as morphological changes in disorders such as MDD.

## Acknowledgements

Authors thank Mehrab Ali and Netta Ussyshkin for support with administrative tasks throughout the study. They thank CAMH animal facility staff for the caring for our animals over the study duration. Authors further thank the members from Charles River Laboratories and NeuroDigitech for their contribution to data generation. We also thank Milwaukee Institute for Drug Discovery and University of Wisconsin-Milwaukee’s Shimadzu Laboratory for Advanced and Applied Analytical Chemistry for help with spectroscopy and National Science Foundation, Division of Chemistry [CHE-1625735].

## Author Contributions

TP and ES designed the study. YM, ZAK, DS and JMC synthesized the compounds tested in this study. TP and PL performed the behavioral piece. AK and MS performed the pharmacokinetic piece. Electrophysiology was outsourced to Charles River Laboratories. TP, AB and MM analyzed the data. TP and AB wrote the manuscript.

