## Supplementary for "Symptomatic and Neurotrophic Effects of GABAA Receptor Positive Allosteric Modulation in a Mouse Model of Chronic Stress"

### Contents

**--- Supplementary Methods ---** 4
Electrophysiology 5
Liver microsomal Assay 5
GL-I-54 Pharmacokinetic Characterization 5
In vitro hydrolytic stability of GL-I-54 in plasma 6
Plasma protein and brain tissue binding of GL-I-54 6
Drug Preparation 6
Chronic Restraint Stress (CRS) 6
UCMS and Randomized Stressors 6
Behavioral Tests 7
Elevated Plus Maze (EPM) 7
Open Field (OF) 7
Novelty Suppressed Feeding (NSF) 7
Phenotyper test 8
Sucrose Consumption (SC) 8
Forced Swim Test (FST) 8
Y Maze 8
Golgi Staining Analysis 9
Statistics 9
**--- Supplementary Tables ---** 11
**Supplementary Table 1.** Statistical significance of potentiation between the different α-subunits tested in presence of GL-I-54 (A) or GL-II-73 (B). 12
**Supplementary Table 2.** Statistical significance of GL-I-54 and GL-II-73 activity on behavioral outcomes (from Figure 1). 13
**Supplementary Table 3.** Statistical significance of acute GL-RM activity on behavioral outcomes (from Figure 2). 14
**Supplementary Table 4.** Statistical significance of chronic GL-RM activity on behavioral outcomes (related to Figure 3) and morphology changes (related to Figure 4). 16
**Supplementary Table 5.** Summary of the correlations between behavioral measures and morphological features after chronic treatment with GL-RM. 19
**--- Supplementary Figures ---** 20

### --- Supplementary Methods ---

#### Electrophysiology

This experiment was outsourced to Charles River Laboratories. Electrophysiological recordings were performed using IonWorks Barracuda™ [1,2]. Full-length cDNAs for the human GABAA receptor subtypes ($\alpha1\beta3\gamma2$, $\alpha2\beta3\gamma2$, $\alpha3\beta3\gamma2$, $\alpha4\beta3\gamma2$ or $\alpha5\beta3\gamma2$) were transfected in HEK-293T cell line (Charles River Laboratories, Cleveland, US). Intracellular solution (50mM CsCl, 90mM CsF, 5mM $MgCl_2$, 1mM EGTA, 10mM HEPES, pH adjusted to 7.2 with CsOH) was loaded into the intracellular compartment of a PPC planar electrode prior to recording. 11μL of extracellular solution, HB-PS (137mM NaCl, 4mM KCl, 3.9mM $CaCl_2$, 1mM $MgCl_2$, 10mM HEPES, 10mM Glucose; pH adjusted to 7.4) was loaded into each well of a PPC plate and cell suspension was added to each well (9μL) of the PPC planer electrode. Whole cell recordings acquired using IonWorks System Software (Molecular Devices Corporation, Union City, CA) established using patch perforation and membrane currents were recorded by on-board patch clamp amplifiers. Holding potential was -70mV. Current recordings after first application of the test articles alone were taken to detect potential agonist effects followed by exposure of each compound in the presence of GABA (Supplementary material Figure 1). During testing, 20μL of test article (GL-I-54 or GL-II-73) was applied to cells (10μL/s) at a range of concentrations (0.033μM, 0.1μM, 0.33μM, 1μM, 3.33μM, 10μM, 33.33μM) and activation of receptors with 5μM GABA. Currents were then normalized to the current elicited by 5μM GABA in the absence of any test article. Of these concentrations, 0.33μM is of physiological relevance and therefore was investigated further. To confirm activity was dependent on the presence of GABA and thus allosteric modulation, recordings were completed in the absence of GABA (Supplemental Figure 2.). From the PubMed Protein Sequence repository for the GABAA receptors subunits, the binding pocket formed by α1/2/3/5 and γ2 is 100% conserved between mouse and human.

#### Liver microsomal Assay

This experiment was outsourced to Charles River Laboratories. Using an Applied Biosystems Sciex API-5500 triple quadrupole mass spectrometer with Sounds Analytics ADDA Autosampler and an equipped electrospray ionization source (600$^{o}$C) operated in the positive-ion mode, the compounds were tested for the metabolic stability in C57BL/6 Mouse liver microsomes (pooled/male 20mg protein/mL) (BiolVT Cat.# M00511, Lot# PQB) at a 2μM concentration and 0.5mg/mL of Matrix concentration. The assay was performed similar to methods described by Di et al.[3]. Briefly, assay plates and were warmed to 37$^{o}$C. NADPH solution and compounds were added to assay plates and then the plate was quenched. Microsomes/S9 were quick thawed and added to KPhos buffer kept at 37$^{o}$C. 200μL of warmed microsomes were added to the assay to start reaction and the time was recorded. For T0, duplicate aliquots were transferred to a quench plate and mixed. This was repeated at 30, 60, and 120. After transfer to quench plate, samples were vortexed and centrifuged at 4$^{o}$C, water was added to a sample plate and then the supernatant was also transferred to the sample plate. Finally, the sample plate was sealed and put into a refrigerated LC/MS/MS autosampler. The half-life and percent remaining of the compound at the end of 120 min are presented in the table#. Each time point was performed in duplicate. Calculations for calculated intrinsic hepatic clearance used 45mg microsomes/g liver and 87.5g liver/kg body weight for mouse.

#### GL-I-54 Pharmacokinetic Characterization

Aiming to obtain the respective pharmacokinetic profile, young adult mice (n = 15) were divided into five groups of animals; each group contained three animals and corresponded to predetermined time intervals (5, 20, 60, 180, and 720 min). The mice were treated intraperitoneally (IP) with a cassette containing four imidazobenzodiazepine compounds each dosed at 3 mg/kg in a total volume of 10 ml/kg. At the appropriate time intervals, the blood samples were collected via cardiac puncture of mice anesthetized with ketamine solution (10% Ketamidor, Richter Pharma Ag, Wels, Austria, dosed IP at 100 mg/kg) in heparinized syringes and centrifuged at 800 rcf for 10 min to obtain plasma samples. Mice were decapitated and brains were weighed, homogenized in 1.25 ml of methanol and centrifuged at 3400 rcf for 20 min. Compounds were extracted from plasma and supernatants of brain tissue homogenates by solid phase extraction, using Oasis HLB cartridges (Waters Corporation, Milford, Massachusetts). The procedure of sample preparation and determination of concentration by ultraperformance liquid chromatography–tandem mass spectrometry (UPLC–MS/MS) with Thermo Scientific Accela 600 UPLC system connected to a Thermo Scientific TSQ Quantum Access MAX triple quadrupole mass spectrometer (Thermo Fisher Scientific, San Jose, California), equipped with electrospray ionization (ESI) source, has been already described in detail [Obradović A. et al. SH-I-048A, an in vitro non-selective super-agonist at the benzodiazepine site of GABAA receptors: the approximated activation of receptor subtypes may explain behavioral effects. Brain Res. 2014 Mar 20;1554:36-48]. Non-compartmental pharmacokinetic analysis was performed using PK Functions for Microsoft Excel software (by Joel Usansky, Atul Desai, and Diane Tang-Liuwere), while graphs were constructed in commercial statistical software Sigma Plot 12 (Systat Software Inc., USA).

#### In vitro hydrolytic stability of GL-I-54 in plasma

The GL-I-54 was tested for its hydrolytic stability in vitro at 37 °C, utilizing blank mouse plasma spiked with the compound and internal standard (SH-I-048A; synthesized at the Department of Chemistry and Biochemistry, University of Wisconsin–Milwaukee, USA), as detailed in: Stamenić TT, Poe MM, Rehman S, Santrač A, Divović B, Scholze P, et al. Ester to amide substitution improves selectivity, efficacy and kinetic behavior of a benzodiazepine positive modulator of GABAA receptors containing the α5 subunit. Eur J Pharmacol. 2016 Nov;791:433–43.

#### Plasma protein and brain tissue binding of GL-I-54

The rapid equilibrium dialysis assay used to determine free fraction of GL-I-54 in mouse plasma and brain tissue was the same as described by Obradović et al.[4]. Free concentrations in the brain were calculated by multiplying the obtained total brain concentrations with the appropriate free fractions determined by rapid equilibrium dialysis.

#### Drug Preparation

Behavioral data for single enantiomers used an acute intraperitoneal injection (IP) depending on the behavioral test (EPM and FST: 5mg/kg, Y maze: 1, 5 and 10mg/kg). GL-I-54 or GL-II-73 was dissolved in vehicle solution (85% distilled $H_2O$, 14% propylene glycol (Sigma Aldrich) and 1% Tween 80 (Sigma Aldrich)).

For the 2 UCMS studies, a racemic preparation containing 50% of GL-I-54 and 50% of GL-II-73 was synthesized, referred to as GL-RM. For acute use, GL-RM was dissolved in vehicle solution, as described above, at a dose of 10mg/kg (effectively 5mg/kg GL-I-54 and 5mg/kg GL-II-73). Solutions were made daily for acute administration. The chronic administration study used the per oral (PO) route with 266mL/kg as the administration volume (calculated from average intake observed within cohorts) amounting to a 30mg/kg dose (effectively 15mg/kg GL-I-54 and 15mg/kg GL-II-73). For this preparation, GL-RM was dissolved in tap water overnight at room temperature and prepared every other day. The GL-RM was then made available to the animals through their drinking water.

#### Chronic Restraint Stress (CRS)

Chronic restraint stress (CRS) was performed in cohorts #2 and #3. CRS involves placing each mouse in a 50mL Falcon™ Tube that is perforated at the tip and lid for airflow. The mouse while inside the tube was returned to home cage for one hour. This was completed twice daily for one week.

#### UCMS and Randomized Stressors

The UCMS paradigm used a variety of mild stressors (1-2 applied daily) that were randomly performed over 6 weeks (8 weeks considering the 2 weeks of behavioral testing). During the testing phase, 1 stressor per day was applied after testing. Animals subjected to UCMS were housed in a separate room without environmental enrichment to exacerbate the effects of other stressors. The following stressors were used randomly during the 6-weeks of UCMS exposure:

1) Dark/light modifications: This stressor perturbs the dark/light cycle of animals with facility room lights being turned on and off at irregular intervals. This stressor was utilized over weekends and never 24 hours prior to testing. Light settings used ranged from 30 min – 4 hours and Dark settings ranged from 3-9 hours.
2) Cage Tilt: Cages are tilted on an approximately $45^o$ angle lengthwise using a PVC pipe and one end for 1-4 hours. During cage tilt, access to food and water is unaffected.
3) Predator Odor: A perforated Falcon™ tube containing a cotton ball exposed to predator urine (Fox (PredatorPee.com)) was placed in the cage of an animal for 1-4 hours.
4) Reduced Space: A rat cage was divided into 4 quadrants using plastic dividers and one mouse is placed in each quadrant. Each mouse has access to food and water during this stressor and was implemented four hours or overnight.

5) Reduced space with odor: 2-3 drops of predator urine (fox) was put in a PVC pipe (9cm diameter, 30cm length) with a plastic bottom and ventilation cap. Mice were then placed in the tubes for 1-4 hours before being returned to their home cage.
6) Restraint: Mice were placed in 50mL Falcon™ tubes with a hole at the tip of the conical bottom and a hole in the lid allowing for adequate ventilation and tail space. Tubes containing the mice were then placed back into respective home cages for 1 hour.
7) Wet Bedding: ~200mL of room temperature water was poured into each cage to wet the bedding. Flooding was avoided as no unabsorbed water was present. Wet bedding stressor was maintained for 1-4 hours and followed by the new bedding stressor.
8) New Bedding: Mice were placed into a fresh newly prepared cage with new bedding. This stressor was avoided 24 hours before testing.
9) No Bedding: All bedding was removed from all cages for 1-3 hours and replaced with new of a mixture of bedding from all cages of the same treatment group.
10) Exchange mice: Mice were removed from their home cage and placed in the home cage of a mouse receiving the same treatment (vehicle or PAM). Time spent in the unfamiliar cage was sustained for hours or overnight.
11) Forced Bath: Bedding was removed from the cage and water was added until filled to a 2-3cm depth from the bottom. After 1 hour, water was removed and a mixture of bedding from animals of the same treatment group was placed in the cages.

#### Behavioral Tests

Prior to behavioral testing, animals were moved to the testing room and allowed to habituate to the room and lighting conditions for 30 minutes. Prior to placing the animal in any apparatus, the apparatus was cleaned with 70% ethanol, and left for 2 minutes to dry in order to remove the odor from the previously tested animal.

#### Elevated Plus Maze (EPM)

The EPM is commonly used in preclinical rodent models to assess anxiety-like behaviors and was used as per standard method[5]. Briefly, mice are placed in a cross-shaped maze raised at 55cm from the floor, with 2 open arms facing each other, and 2 closed arms facing each other. During testing, mice were individually placed in the center of the maze facing an open arm and allowed to explore for 10 minutes. Throughout testing, the exploration of the each mouse was recorded using a digital camera mounted on the ceiling. After 10 minutes of exploration the camera was stopped and the mouse was removed from the maze and returned to its home cage. The videos were analyzed using Ethovision XT14 software. Specific parameters considered were: % time spent in the open arms, calculated using ([(total time spent in open arms)/(total time spent in open arms + total time spent in closed arms)]*100) and % open arm entries calculated using ([(total open arm entries)/(total open arm entries + total closed arm entries)]*100).

#### Open Field (OF)

The OF test is designed to test anxiety-like behavior in rodents through their ambulatory movements in an enclosed arena. The open field test was run in a 43x43x50cm square chamber with grey floor and dark grey walls. One mouse was placed in a single field and recorded for 10 minutes by the same system used in the EPM. Time spent and number of entries into the center were used as measures of anxiety-like behaviors.

#### Novelty Suppressed Feeding (NSF)

NSF is used to detect stress-induced anxiety-like behavior by measuring a rodent's reluctance to eat in an unfamiliar environment. The NSF test chamber is a 62x31x40cm Plexiglas container with clear floors and walls. Mice were weighed and food-deprived for 16 hours prior to testing. The following day, animal weights were assessed ensuring an 8-12% weight loss. If a mouse did not meet this criteria, it was not tested. A familiar kind of food pellet was placed ~8 cm from the wall of the chamber on the side closest to the experimenter and the mouse was placed on the opposite side of the chamber. A timer was started immediately and the latency to approach and consume the food pellet was recorded by an experimenter. After the test was completed, mice were returned to their home cage where latency to bite/feed on the food pellet was measured to control for appetite drive or food deprivation-induced weight loss. A maximum of 12 minutes is allotted for testing in the novel environment and 6 minutes in the animal's home environment.

#### Phenotyper test

The Noldus™ PhenoTyper (Leesburg, VA, USA) is an apparatus that allows for observation of mice over an extended length of time and in response to various stimuli. The PhenoTyper box consists of a clear tray (30x30cm) surrounded by clear walls. One wall has an opening to allow the experimenter to access the inside of the chamber easily; another wall has a metal food tank attached and a small hole for a water bottle nozzle. A top unit encloses the apparatus on the top and has a built in infrared sensitive camera, infrared LED lights and an audio stimulus. Inside the phenotyper, white corn-bedding was added to the bottom tray (2cm depth) and a shelter was secured to the wall in the back right corner. In brief, the setup is equipped with a shelter and a white light shining above the food tank. The test is performed overnight, during the active phase of the animal. Mice are placed in the PT at 18:00hr (1 mouse/box) and monitored until 8:00hr the next morning. From 18:00hr - 23:00hr a 5 hour baseline is recorded and at 23:00 a 1 hour light challenge is initiated. From 23:00hr - 0:00hr a light shines directly on the food tank as an aversive stimulus as this is the time-window when mice naturally have a high plateau for time within the food zone. From 0:00hr - 8:00hr behavioral patterns and coping behaviors are observed. Activity of the animal is recorded via a camera mounted in the ceiling of the box, and the activity is measured using the tracking system Ethovision (Noldus). Mice can hide in a shelter, and the time spent in the shelter is measured overnight. The residual avoidance score captures the time spent in the shelter, after the light is turned back off, measuring lasting avoidance of the previously lit zone. The design of this assay includes characterization of the activity of the mouse before, during and after the stressful stimulus.

After a baseline characterization of the animal behavior, mice are split into 2 groups, control or UCMS. During the UCMS protocol, mice were tested in the PT boxes each week in addition. For the acute drug treatment study, mice were acutely injected with saline or GL-RM (10mg/kg IP) before 20:00hr (Depicted by the black arrow in Figure 2f). Anxiety-like behavior can be reproducibly quantified by investigating time spent in shelter zone (using 1 hour bins) and residual avoidance can be calculated. Residual avoidance (RA) is an index calculated as a function of the control group response to the light challenge and subsequent 5 hours, as per method described in Prevot et al (2019). A positive RA value indicates avoidance of the lit zone and is calculated using the following equations based on the time spent in the shelter zone. Shelter Zone: [(Σ Time (12am–5am)–Time (11pm–12am))/Average control group (Σ Time (12am–5am)-Time (11pm–12am))-1]*100, where "Time" is time spent in shelter zone. Chronically stressed mice have a predictable avoidance behavioral pattern after an aversive light stimulus that is detectable in the PhenoTyper™ [5].

#### Sucrose Consumption (SC)

Sucrose consumption is used to measure anhedonia-like behavior. Mice were habituated to a 1% sucrose drinking solution for 48 hours followed by fluid-deprivation overnight (~16 hours). To test sucrose consumption, the weight of 1% sucrose solution consumed from a tube given to individually-housed mice is measured for 1 hour. Similar design is used for water consumption the following day. % Sucrose consumption was then calculated using the following equation: % Sucrose Consumption = [(Sucrose consumed)/sucrose consumed + water consumed)]*100. If an animal did not drink any sucrose they were not included in the analysis as they cannot show preference.

#### Forced Swim Test (FST)

FST is used to detect antidepressant-like activity of compounds. Mice were placed in a 4 liter beaker (1 mouse /beaker) containing ~3 liters of water (23$^{o}$C-25$^{o}$C) for a total of 6 minutes. A digital camera filmed from the side. After 6 minutes, mice were removed from the water, dried with a towel and warmed under a heat lamp for 5 minutes. Videos were analyzed using EthoVision XT 14 software after acquisition. Total time immobile (immobility defined as the minimum movement to keep the head above water) was analyzed for minutes 0-4 and reported.

#### Y Maze

The Y maze is a spatial working memory paradigm looking at spontaneous alternation and adapted from the T maze paradigm described by Faucher et al[6]. The Maze has 3 arms separated at 120$^{o}$ from each other, each 26cm long, 8cm wide and 13 cm high. Two arms ("goal arms") have a sliding door close to the center of the maze and one arm ("start box") has a sliding door close to the distal part of the arm. This arm is the arm closest to the experimenter and where a mouse is placed to begin each trial. A habituation phase is conducted first where mice are habituated to the maze over 2 consecutive days. In this phase each animal is allowed to explore the maze freely for 10 minutes. On the third day a training phase is initiated. This consists of seven successive trials separated by a 30 second inter-trial interval (ITI). During this training the animal is placed in the start box for 30 seconds, the sliding door is then opened and the mouse can enter either of the goal arms. Once the goal arm is chosen, that arm door is closed and the choice and latency to decide are recorded. The animal is left in the chosen arm for 30 seconds before being returned to the start box for the next trial, which starts with the initial confinement in the start box for 30 seconds. This is repeated over 7 trials. Between each trial the maze is wiped down lightly with 70% ethanol to remove olfactory cues. The same procedure used for the training phase was used for the testing phase (24 hours later) however the ITI was increased to 90 seconds. Also, on the 8$^{th}$ trial the above process is repeated however the inter-trial interval is reduced to 10 seconds to test the animal's motivation to alternate. During the testing phase, animals that do not alternate on the 8$^{th}$ trial show reduced motivation and therefore are excluded form analysis as their results are biased by lack of motivation and not cognitive performance. The mean alternation rate was calculated using the following formula and expressed as a percent: Mean alternation % = [(Number of alternations/number of trials)*100].

#### Golgi Staining Analysis

Neuronal morphology was assessed using Golgi Staining. The analysis was performed by NeuroDigiTech (San Francisco CA., USA). After arrival, a cryostat was used to cut 100µm thick serial, coronal slices from anterior to posterior and slices were mounted on glass slides. Basal and apical dendrites of pyramidal cells were identified in layers II/III of the PFC and in the CA1 of the hippocampus and determined to be the regions of interest (ROIs). For quantification, NeuroLucida v10 (Microbrightfield, VT) software was used on a Dell PC computer and controlled a Zeiss Axioplan 2 image microscope with Optronics MicroFire CCD camera (1600x1200). A motorized X, Y and Z focus allowed for high resolution images and later quantification. ROIs were first considered under low magnification (10x and 20x) to identify those with the least truncations of distal dendrites at high-magnification (40x and 60x). Zeiss 100x objective was then used with immersion oil to construct 3D images and allow for spine counting along the complete dendritic tree. Neurons analyzed (n=6 per animal) were required to visually have a completely filled soma and dendrites, not overlap with other soma and have complete 3D visualization of the dendritic tree. Neurons with incomplete staining and/or incomplete visualization due to section plane were not included in the analysis. Analysis protocol and inclusion/exclusion criteria were adapted from Wu et al.[7]. For spine sampling, only those orthogonal to the dendritic shaft were well characterized and included as those above or below could not be adequately distinguished. Raw data was extrapolated and quantified using NeuroExplorer (Microbrightfield, VT).

#### Statistics

##### *Z-Scores Calculation*

Z-scores were calculated to provide an overall measure of anxiety-like and depressive-like behavioral phenotypes, referred as z-emotionality[8]. Z-scores for each animal were calculated for each parameter of the behavioral tests using the following formula: Z score= [(Individual raw data-Average control raw data)/(Standard deviation control raw data)] or Z score= [(Individual raw data-Average control raw data)/-(Standard deviation control raw data)] dependent on the original test's raw values (higher values for higher anxiety, or lower values for higher anxiety, respectively). A combined behavioral emotionality Z-scores was then calculated for each animal, derived from EPM Z-scores (%Time spent in open arms and % open arm entries), NSF (Latency to approach and latency to bite), PhenoTyper (week 6 food zone residual avoidance and shelter zone residual avoidance), coat state, FST (time immobile) and sucrose consumption at week 6. In the chronic GL-RM study one outlier was removed in the Control with GL-RM group.

##### *Statistical Testing*

Statistical analyses were performed using GraphPad Prism 9 software. Electrophysiology data of GL-II-73 and GL-I-54 and their activity at GABAA receptor subunits was compared to 100% using T-test. Then, overall activity was analyzed using two-way ANOVA. Independent variables were considered as concentration and receptor subunit type. Bonferroni post-hoc testing was used to identify differences at specific subunits. GL-I-54 EPM and FST behavioral testing were analyzed using unpaired t-tests and Y maze testing was analyzed using one-way ANOVA with Fisher's lest significant difference post-hoc test (PLSD). UCMS studies that involved weekly assessment were analyzed using repeated measures two-way ANOVA (coat state, weight gain, PhentoTyper time spent in specific zones and residual avoidance scores). Standard two-way ANOVA was used to evaluate all other behavioral outcomes from the different tests (EPM, OF, NSF, SC, FST and Y maze) considering both UCMS and treatment as independent variables. Fishers PLSD tests were used for post hoc analysis. Golgi staining employed repeated measures one-way ANOVA with Tukeys post hoc tests. All correlation analyses were conducted using Spearman ranks correlation between morphological features and behavioral outcomes. For all comparisons, statistical significance was considered $p<0.05$ and all data are presented as the mean ± SEM.

### --- Supplementary Tables ---

***Supplementary Table 1.*** *Statistical significance of potentiation between the different α-subunits tested in presence of GL-I-54 (A) or GL-II-73 (B).*

| n=4 | | GL-I-54 | | | | | | | | | |
|---|---|---|---|---|---|---|---|---|---|---|---|
| | | α1vα2 | α1vα3 | α1vα4 | α1vα5 | α2vα3 | α2vα4 | α2vα5 | α3vα4 | α3vα5 | α4vα5 |
| Concentration (µM) | 0.03 | ns | ns | ns | ns | ns | ns | ns | ns | ns | ns |
| | 0.1 | ns | **$p<0.05$** | ns | ns | ns | ns | ns | **$p<0.05$** | ns | ns |
| | 0.33 | **$p<0.01$** | **$p<0.001$** | ns | ns | ns | **$p<0.001$** | ns | **$p<0.001$** | ns | ns |
| | 1 | ns | **$p<0.001$** | ns | ns | ns | **$p<0.001$** | ns | **$p<0.001$** | **$p<0.05$** | **$p<0.01$** |
| | 3.33 | ns | **$p<0.001$** | **$p<0.01$** | ns | **$p<0.001$** | **$p<0.001$** | **$p<0.01$** | **$p<0.001$** | **$p<0.001$** | ns |
| | 10 | ns | **$p<0.001$** | **$p<0.01$** | ns | **$p<0.001$** | **$p<0.001$** | ns | **$p<0.001$** | **$p<0.001$** | **$p<0.01$** |
| | 33.3 | ns | **$p<0.001$** | **$p<0.01$** | ns | **$p<0.001$** | **$p<0.001$** | **$p<0.05$** | **$p<0.001$** | **$p<0.001$** | ns |
| n=4 | | GL-II-73 | | | | | | | | | |
| Concentration (µM) | 0.03 | ns | ns | ns | **ns** | ns | ns | **ns** | ns | **ns** | **ns** |
| | 0.1 | ns | ns | ns | **ns** | ns | ns | **ns** | ns | **ns** | **ns** |
| | 0.33 | ns | ns | ns | **$p<0.05$** | ns | ns | **ns** | ns | **ns** | **ns** |
| | 1 | ns | ns | ns | **$p<0.05$** | ns | ns | **ns** | ns | **$p<0.05$** | **ns** |
| | 3.33 | ns | ns | ns | **$p<0.001$** | ns | ns | **$p<0.01$** | ns | **$p<0.01$** | **$p<0.001$** |
| | 10 | ns | ns | ns | **$p<0.001$** | ns | ns | **$p<0.001$** | ns | **$p<0.001$** | **$p<0.001$** |
| | 33.3 | **$p<0.05$** | **$p<0.05$** | ns | **$p<0.001$** | ns | **$p<0.001$** | **ns** | **$p<0.001$** | **ns** | **$p<0.001$** |

***Supplementary Table 2.*** *Statistical significance of GL-I-54 and GL-II-73 activity on behavioral outcomes (from Figure 1).*

| Assay | Statistical test | F-value | p-value | Sex Effect |
|---|---|---|---|---|
| **EPM** | One-way ANOVA | $F_{(3,20)}=1.563$ | p=0.2296 | p>0.05 |
| **FST** | One-way ANOVA | $F_{(3,20)}=8.264$ | **p=0.0009*** ** | p>0.05 |
| | *PLSD (Control vehicle vs CRS vehicle* | | **p=0.0265*** | |
| | *PLSD (Control vehicle vs GL-I-54)* | | **p=0.0272*** | |
| | *PLSD (CRS vehicle vs CRS GL-I-54* | | **p=0.0001*** ** | |
| | *PLSD (CRS GL-I-54 vs CRS GL-II-73* | | **p=0.0021** ** | |
| **Y Maze** | One-way ANOVA | $F_{(4,29)}=6.68$ | **p=0.0006*** ** | p>0.05 |
| | *PLSD (Control-Vehicle vs CRS-Vehicle)* | | **p<0.001*** ** | |
| | *PLSD (CRS-10mg/kg vs CRS-Vehicle)* | | **p=0.02*** | |
| **Rotarod** | Repeated Measure ANOVA | $F_{(1,18)}=6.68$ | **p=0.0024** ** | p>0.05 |
| | *PLSD (Control-Vehicle vs Gl-I-54 10mg/kg; 5 min)* | | p=0.1 | |
| | *PLSD (Control-Vehicle vs Gl-I-54 10mg/kg; 20 min)* | | p=0.1 | |
| | *PLSD (Control-Vehicle vs Gl-I-54 10mg/kg; 60min)* | | **p=0.02*** | |

***Supplementary Table 3.*** *Statistical significance of acute GL-RM activity on behavioral outcomes (from Figure 2).*

| Assay | Statistical test | F-value | p-value | Sex Effect |
|---|---|---|---|---|
| **EPM (% time in open arms)** | Two-way ANOVA<br>UCMS<br>Treatment<br>Interaction | <br>$F_{(1;44)}$=2.33<br>$F_{(1;44)}$=0.1397<br>$F_{(1;44)}$=1.122 | <br>p=0.13<br>p=0.71<br>p=0.30 | p>0.05 |
| **EPM (% entries in open arms)** | Two-way ANOVA<br>UCMS<br>Treatment<br>Interaction | <br>$F_{(1;44)}$=3.47<br>$F_{(1;44)}$=0.03<br>$F_{(1;44)}$=0.60 | <br>p=0.07[t]<br>p=0.87<br>p=0.44 | p>0.05 |
| **NSF (Latency to approach)** | Two-way ANOVA<br>UCMS<br>Treatment<br>Interaction | <br>$F_{(1;41)}$=2.93<br>$F_{(1;41)}$=0.017<br>$F_{(1;41)}$=5.48 | <br>p=0.09[t]<br>p=0.90<br>**p=0.02*** | p>0.05 |
| | *PLSD (Control vs UCMS)* | | <br>**p=0.011*** | |
| | *PLSD (Vehicle vs GL-RM)* | | **p=0.02*** | |
| **NSF (Latency to bite)** | Two-way ANOVA<br>UCMS<br>Treatment<br>Interaction | <br>$F_{(1;41)}$= 0.67<br>$F_{(1;41)}$= 0.59<br>$F_{(1;41)}$= 0.98 | <br>p=0.42<br>p=0.45<br>p=0.33 | p>0.05 |
| **Phenotyper (time in shelter on week 6)** | Repeated measure ANOVA<br>UCMS<br>Treatment<br>Interaction | <br><br>$F_{(1;44)}$=37.72<br>$F_{(1;44)}$=1.52<br>$F_{(1;44)}$=0.2 | <br><br>**p=<0.001*****<br>p=0.22<br>p=0.66 | **p=0.007**** |
| | *Bonferroni (Control vehicle vs UCMS vehicle)*<br>*10pm*<br>*11pm*<br>*12am*<br>*1am* | | <br><br><br>**p<0.001*****<br>**p<0.01****<br>**p<0.001*****<br>**p<0.001***** | |

| Assay | Statistical test | F-value | p-value | Sex Effect |
|---|---|---|---|---|
| **Phenotyper (Residual Avoidance)** | Two-way ANOVA<br>UCMS<br>Treatment<br>Interaction | <br>$F_{(1;44)}$=23.964<br>$F_{(1;44)}$=0.18<br>$F_{(1;44)}$=0.5 | <br>**p<0.0001*****<br>p=0.67<br>p=0.48 | **p=0.0002***** |
| | *PLSD (Control vs UCMS)* | | **p=<0.0001***** | |
| | *PLSD (Vehicle vs GL-RM)* | | p=0.63 | |
| **Sucrose Consumption** | Two-way ANOVA<br>UCMS<br>Treatment<br>Interaction | <br>$F_{(1;42)}$=4.23<br>$F_{(1;42)}$=0.26<br>$F_{(1;42)}$=1.97 | <br>**p=0.046***<br>p=0.62<br>p=0.17 | p>0.05 |
| **FST†** | Two-way ANOVA<br>UCMS<br>Treatment<br>Interaction | <br>$F_{(1;43)}$=12.48<br>$F_{(1;43)}$=7.611<br>$F_{(1;43)}$=0.009 | <br>**p=0.001*****<br>**p=0.008****<br>p=0.92 | p>0.05 |
| **Z-Score Emotionality** | Two-way ANOVA<br>UCMS<br>Treatment<br>Interaction | <br>$F_{(1;40)}$=47.83<br>$F_{(1;40)}$=2.2<br>$F_{(1;40)}$=0.30 | <br>**p<0.0001*****<br>p=0.15<br>p=0.58 | p>0.05 |
| **Y Maze** | Two-way ANOVA<br>UCMS<br>Treatment<br>Interaction | <br>$F_{(1;18)}$=11.62<br>$F_{(1;18)}$=1.75<br>$F_{(1;18)}$=16.85 | <br>**p=0.003****<br>p=0.20<br>**p=0.0007***** | p>0.05 |
| | *PLSD (Control-Vehicle vs UCMS-Vehicle)* | | **p=0.0013**** | |
| | *PLSD (UCMS-Vehicle vs UCMS-GL-RM)* | | **p=0.0022**** | |

***Supplementary Table 4.*** *Statistical significance of chronic GL-RM activity on behavioral outcomes (related to Figure 3) and morphology changes (related to Figure 4).*

| Assay | Statistical test | F-value | p-value | Sex Effect |
|---|---|---|---|---|
| **EPM (% time in open arms)** | Two-way ANOVA | | | p>0.05 |
| | UCMS | $F_{(1;44)}=0.56$ | p=0.46 | |
| | Treatment | $F_{(1;44)}=0.51$ | p=0.48 | |
| | Interaction | $F_{(1;44)}=0.50$ | p=0.48 | |
| **EPM (% entries in open arms)** | Two-way ANOVA | | | p>0.05 |
| | UCMS | $F_{(1;44)}=0.1197$ | p=0.73 | |
| | Treatment | $F_{(1;44)}=0.0774$ | p=0.78 | |
| | Interaction | $F_{(1;44)}=2.934$ | $p=0.09^{t}$ | |
| **OF (time in center)** | Two-way ANOVA | | | p>0.05 |
| | UCMS | $F_{(1;44)}=0.05$ | p=0.82 | |
| | Treatment | $F_{(1;44)}=0.71$ | p=0.41 | |
| | Interaction | $F_{(1;44)}=1.05$ | p=0.31 | |
| **OF (entries in center)** | Two-way ANOVA | | | p>0.05 |
| | UCMS | $F_{(1;44)}=0.47$ | p=0.5 | |
| | Treatment | $F_{(1;44)}=3.49$ | $p=0.06^{t}$ | |
| | Interaction | $F_{(1;44)}=0.78$ | p=0.38 | |
| **NSF (latency to approach)** | Two-way ANOVA | | | p>0.05 |
| | UCMS | $F_{(1;44)}=0.5$ | p=0.49 | |
| | Treatment | $F_{(1;44)}=0.41$ | p=0.53 | |
| | Interaction | $F_{(1;44)}=0.54$ | p=0.47 | |
| **NSF (latency to bite)** | Two-way ANOVA | | | p>0.05 |
| | UCMS | $F_{(1;44)}=6.9$ | **p=0.01**** | |
| | Treatment | $F_{(1;44)}=6.7$ | **p=0.01**** | |
| | Interaction | $F_{(1;44)}=1.26$ | p=0.27 | |
| **Phenotyper (time in shelter on week 6)** | Repeated measure ANOVA | | | **p=0.0001***** |
| | UCMS | $F_{(1;44)}=5.03$ | **p=0.03*** | |
| | Treatment | $F_{(1;44)}=0.009$ | p=0.92 | |
| | Interaction | $F_{(1;44)}=2.28$ | p=0.14 | |

| Assay | Statistical test | F-value | p-value | Sex Effect |
|---|---|---|---|---|
| | *Bonferroni (Control vehicle vs UCMS vehicle)*<br>*12am*<br>*1am* | | <br><br>**p<0.05***<br>**p<0.01**** | |
| **Phenotyper (Residual avoidance)** | Repeated measure ANOVA<br>UCMS<br>Treatment<br>Interaction | <br>$F_{(1;44)}$=60.08<br>$F_{(1;44)}$= 2.82<br>$F_{(1;44)}$=2.38 | <br>**p<0.001*****<br>p=0.09[t]<br>p=0.13 | p>0.05 |
| | *Bonferroni (Control vehicle vs UCMS vehicle)*<br>*Week 3*<br>*Week 4*<br>*Week 5*<br>*Week 6* | | <br>**p<0.001*****<br>**p<0.001*****<br>**p<0.01****<br>**p<0.001***** | |
| **Sucrose Consumption** | Two-way ANOVA<br>UCMS<br>Treatment<br>Interaction | <br>$F_{(1;44)}$=1.6<br>$F_{(1;44)}$=6.2<br>$F_{(1;44)}$=0.07 | <br>p=0.21<br>**p=0.016**<br>p=0.8 | p>0.05 |
| **FST** | Two-way ANOVA<br>UCMS<br>Treatment<br>Interaction | <br>$F_{(1;44)}$=0.03<br>$F_{(1;44)}$=0.03<br>$F_{(1;44)}$=1.21 | <br>p=0.88<br>p=0.85<br>p=0.28 | p>0.05 |
| **Z-Score Emotionality[†]** | Two-way ANOVA<br>UCMS<br>Treatment<br>Interaction | <br>$F_{(1;43)}$=5.38<br>$F_{(1;43)}$=3.09<br>$F_{(1;43)}$=0.002 | <br>**p=0.03***<br>**p=0.08[t]**<br>p=0.97 | p>0.05 |
| **Y Maze** | Two-way ANOVA<br>UCMS<br>Treatment<br>Interaction | <br>$F_{(1;37)}$=6.33<br>$F_{(1;37)}$=9.26<br>$F_{(1;37)}$=4.76 | <br>**p=0.016***<br>**p=0.0043****<br>**p=0.03*** | p>0.05 |
| | *PLSD (Control-Vehicle vs UCMS-Vehicle)* | | **p=0.005**** | |

| Assay | Statistical test | F-value | p-value | Sex Effect |
|---|---|---|---|---|
| | *PLSD (UCMS Vehicle vs UCMS-GL-RM)* | | **p=0.0006*** ** | |
| **Golgi (spine count-basal) - PFC** | Repeated measure ANOVA | $F_{(2;69)}$=25.28 | **p<0.0001*** ** | p>0.05 |
| | *Tukeys' (control versus UCMS)* | | **p<0.0001*** ** | |
| | *Tukeys' (UCMS versus UCMS+GLRM)* | | **p<0.0001*** ** | |
| | *Tukeys' (Control versus UCMS+GLRM)* | | **p=0.0128* ** | |
| **Golgi (spine count-apical) - PFC** | Repeated measure ANOVA | $F_{(2;69)}$=26.01 | **p<0.0001*** ** | p>0.05 |
| | *Tukeys' (control versus UCMS)* | | **p<0.0001*** ** | |
| | *Tukeys' (UCMS versus UCMS+GLRM)* | | **p<0.0001*** ** | |
| | *Tukeys' (Control versus UCMS+GLRM)* | | p=0.67 | |
| **Golgi (spine count-basal) – CA1** | Repeated measure ANOVA | $F_{(2;69)}$= 34.56 | **p<0.0001*** ** | p>0.05 |
| | *Tukeys' (control versus UCMS)* | | **p<0.0001*** ** | |
| | *Tukeys' (UCMS versus UCMS+GLRM)* | | **p<0.0001*** ** | |
| | *Tukeys' (Control versus UCMS+GLRM)* | | **p<0.001*** ** | |
| **Golgi (spine count-apical) – CA1** | Repeated measure ANOVA | $F_{(2;69)}$= 19.73 | **p<0.0001*** ** | p>0.05 |
| | *Tukeys' (control versus UCMS)* | | **p<0.0001*** ** | |
| | *Tukeys' (UCMS versus UCMS+GLRM)* | | **p<0.01** ** | |
| | *Tukeys' (Control versus UCMS+GLRM)* | | **p<0.001** ** | |
| **† Outlier Removed** | | | | |

***Supplementary Table 5.*** *Summary of the correlations between behavioral measures and morphological features after chronic treatment with GL-RM.*

| Morphological Feature | | | | Behavioral Measure | |
|---|---|---|---|---|---|
| | | | | Y-Maze | Z-Emotionality |
| PFC | Spine Count | Basal | | **Spearman r=0.6916; p=0.0127*** | Spearman r=-0.02098; p=0.9560 |
| | | Apical | | Spearman r=0.3829; p=0.2192 | Spearman r=0.02797; p=0.9388 |
| | Dendritic Length | Basal | | Spearman r=-0.02735; p=0.9328 | Spearman r=0.006993; p=0.9910 |
| | | Apical | | Spearman r=0.05471; p=0.8659 | Spearman r=0.4056; p=0.1928 |
| | Spine Density | Basal | | **Spearman r=0.7616; p=0.004**** | Spearman r=-0.007005; p=0.9865 |
| | | Apical | | **Spearman r=0.6217; p=0.0309*** | Spearman r=-0.2324; p=0.4681 |
| CA1 | Spine Count | Basal | | **Spearman r=0.6486; p=0.0225*** | Spearman r=-0.5175; p=0.0888[t] |
| | | Apical | | **Spearman r=0.7424; p=0.0057**** | Spearman r=-0.4685; p=0.1275 |
| | Dendritic Length | Basal | | Spearman r=0.2892; p=0.3620 | **Spearman r=-0.7622; p=0.0055**** |
| | | Apical | | Spearman r=0.2266; p=0.4787 | **Spearman r=-0.7133; p=0.0118*** |
| | Spine Density | Basal | | **Spearman r=0.6615; p=0.0191*** | Spearman r=-0.4623; p=0.1314 |
| | | Apical | | **Spearman r=0.7496; p=0.0050**** | Spearman r=-0.4343; p=0.1587 |

### --- Supplementary Figures ---

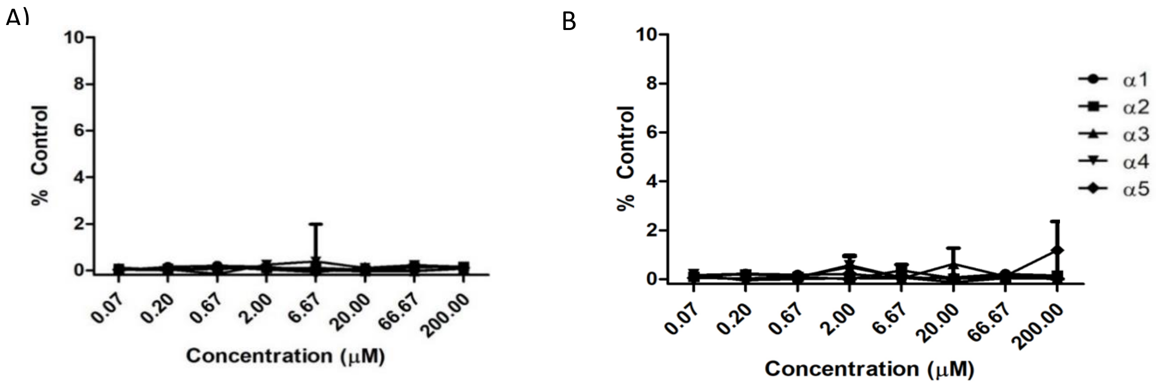

***Supplementary Figure 1.*** *Electrophysiological profile of GL-I-54 and GL-II-73 at α1-5-containing receptors in absence of GABA in the media*

Agonist effect or the lack thereof, GL-I-54 (A) and GL-II-73 (B) were tested in HEK-296 cells expressing recombinant α1-5-GABAA receptors, in absence of GABA in the media, in order to test potential agonist effect. As expected, neither GL-I-54 nor GL-II-73 induced potentiation at any of the receptors, confirming their role as allosteric modulators and not agonists.

| 3 mg/kg | 5 mg/kg (estimation) | | | |
| --- | --- | --- | --- | --- |
| brain concentration | | | | |
| total | | free | | |
| ng/g | | | ng/ml | nmol/l |
| 311.75 | 520.62 | 59.82 | 57.52 | 148.98 |
| 264.61 | 441.90 | 53.65 | 51.59 | 133.62 |
| / | x1.67 | xFF | /1.04 | x2.59 |
| Step 1 | Step 2 | Step 3 | Step 4 | Step 5 |

| % response to GABA alone | | | | | | | |
| --- | --- | --- | --- | --- | --- | --- | --- |
| $\alpha_1$ | | $\alpha_2$ | | $\alpha_3$ | | $\alpha_5$ | |
| 100 | 330 | 100 | 330 | 100 | 330 | 100 | 330 |
| + | + | ++ | +++ | +++ | +++ | +++ | ++ |
| Ø | Ø | + | Ø | Ø | Ø | Ø | ++ |
| Ø < 120% | | + < 200% | | ++ < 250% | | +++ ≥ 250% | |
| Step 6 | | | | | | | |

estimated range of potentiation of $GABA_A$ receptors after 5 mg/kg dose

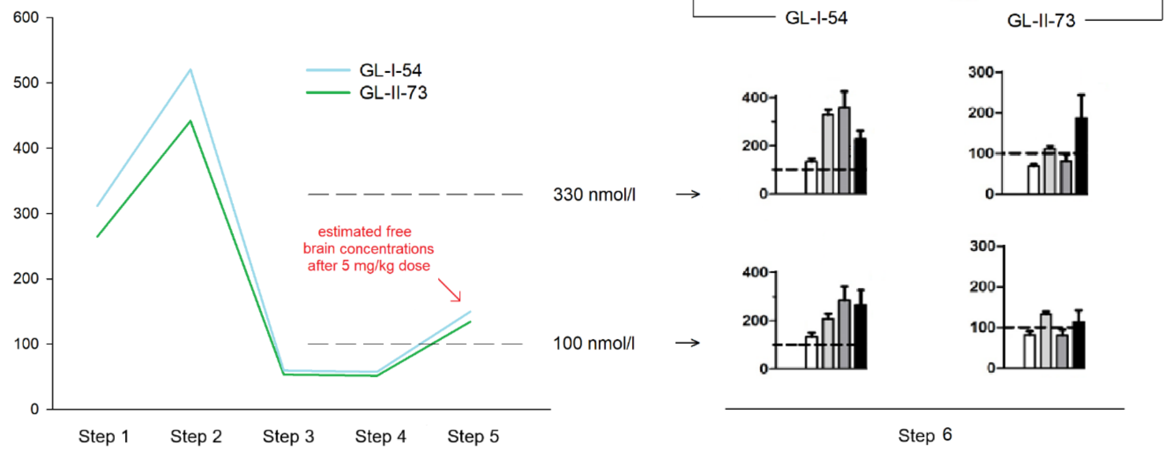

***Supplementary Figure 2.*** *Estimate maximum free brain concentrations of 5 mg/kg of GL-I-54 and 5 mg/kg of GL-II-73*

Stepwise approach to estimate the maximum free brain concentrations of 5 mg/kg of GL-I-54 and 5 mg/kg of GL-II-73 (doses used in acute behavioral testing), and associate them with the range of corresponding $GABA_A$ receptor potentiation values (obtained in electrophysiological experiments). Maximum brain concentrations are achieved around 20 min after dosing (**Figure 1C** and Prevot et al.[9]). Blue content refers to GL-I-54 and green to GL-II-73. White and nuances of the grey and black color at the right bottom graph refer to the particular α subunit of $GABA_A$ receptors, as indicated in the upper right table. **Step 1** – brain concentrations (ng/g) determined 20 min after intraperitoneal dosing of 3 mg/kg GL-I-54 or GL-II-73 to male mice, respectively (n = 3 per compound); **Step 2** – estimation of maximum brain concentrations (ng/g) for 5 mg/kg of GL-I-54 and 5mg/kg of GL-II-73, obtained by multiplying Step 1 concentrations with the factor 5/3 (=1.67), based on an assumption that both GL-I-54 and GL-II-73 display a first-order pharmacokinetics (pharmacokinetics of GL-II-73 is not saturated at 3 and 10 mg/kg, since the increase in the concentration of the compound is proportionate); **Step 3** – free brain concentrations (ng/g) obtained by multiplying Step 2 concentrations with the free fractions in the brain determined by rapid equilibrium dialysis (0.1149 for GL-I-54 and 0.1214 for GL-II-73); **Step 4** – adjusting the Step 3 concentrations (ng/g) for the brain density (1.04 g/ml) and converting the units to ng/ml; **Step 5** – further conversion of units to nmol/l (unit used in electrophysiological experiments) by multiplying Step 4 concentrations with 1000/386.42 (386.42 g/mol is a molecular weight of GL-I-54 and GL-II-73) – the comparable estimated maximum free brain concentrations (nmol/l) of 5 mg/kg GL-I-54 and 5 mg/kg of GL-II-73 are now obtained; **Step 6** – association of Step 5 concentrations with the electrophysiological results – the estimated concentrations closely correspond to 100 nmol/l or, to a lesser extent, to 330 nmol/l of a compound administered in electrophysiological experiment. This implies that, in acute behavioral experiments with 5 mg/kg dose, GL-I-54 likely potentiates $\alpha_1$, $\alpha_2$, $\alpha_3$ and $\alpha_5$ $GABA_A$ receptors, however, $\alpha_1$ potentiation is mild. 5 mg/kg of GL-II-73, on the other hand, is fairly silent on all $GABA_A$ receptors, with the potential to mildly activate $\alpha_2$ and $\alpha_5$ receptors; on $\alpha_1$ and $\alpha_3$ receptors, nevertheless, it behaves as a null modulator. Due to such properties, the racemic mixture of 5 mg/kg of GL-I-54 and 5 mg/kg of GL-II-73 might display advantageous pharmacological profile in vivo. By complete attenuation of activity at α1 $GABA_A$ receptors with GL-II-73, the potentiation of α2, α3 and α5 $GABA_A$ receptors by GL-I-54 might become predominant, exhibiting favorable behavioral effects. Ø – potentiation < 120% (null), + – potentiation < 200% (mild), ++ – potentiation < 250% (moderate), +++ – potentiation ≥ 250 % (strong). For exact values of potentiation at $GABA_A$ receptors containing distinct α subunits, please refer to **Figure 1.**

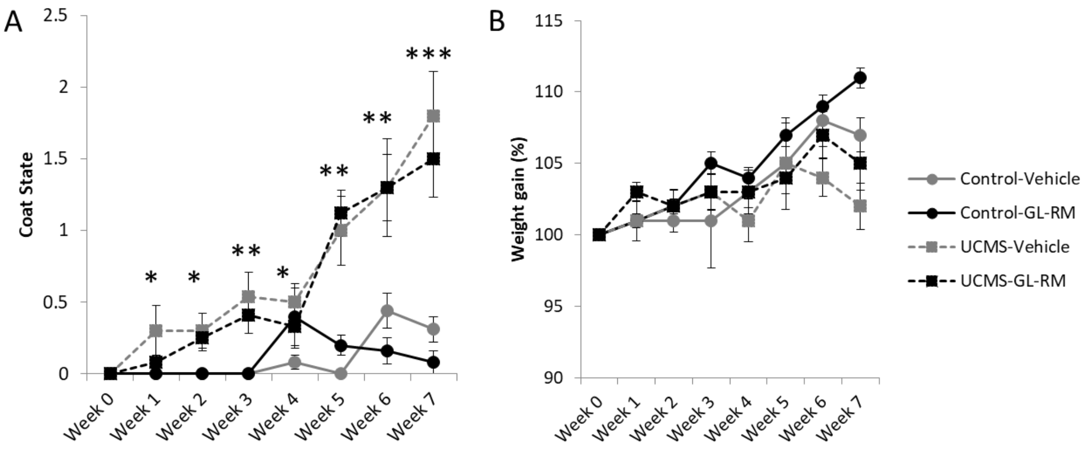

***Supplementary Figure 3.*** *Coat state and Weight Gain changes over the weeks*

Coat state (A) and weight gain (B) were assessed on a weekly basis. Coat state showed a progressive increase in deterioration of the coat state in animals exposed to UCMS ($F_{(1;44)}$=36.437; p<0.0001), regardless of the treatment group (p>0.7807). *Post hoc* testing of UCMS characterized significant coat state deterioration beginning at week 1 (p=0.0228) through until week 7. Weight gain did not show significant changes between groups. There was no sex effect on coat state or weight gain. *p<0.05, **p<0.01, ***p<0.001 compared to Control-Vehicle.

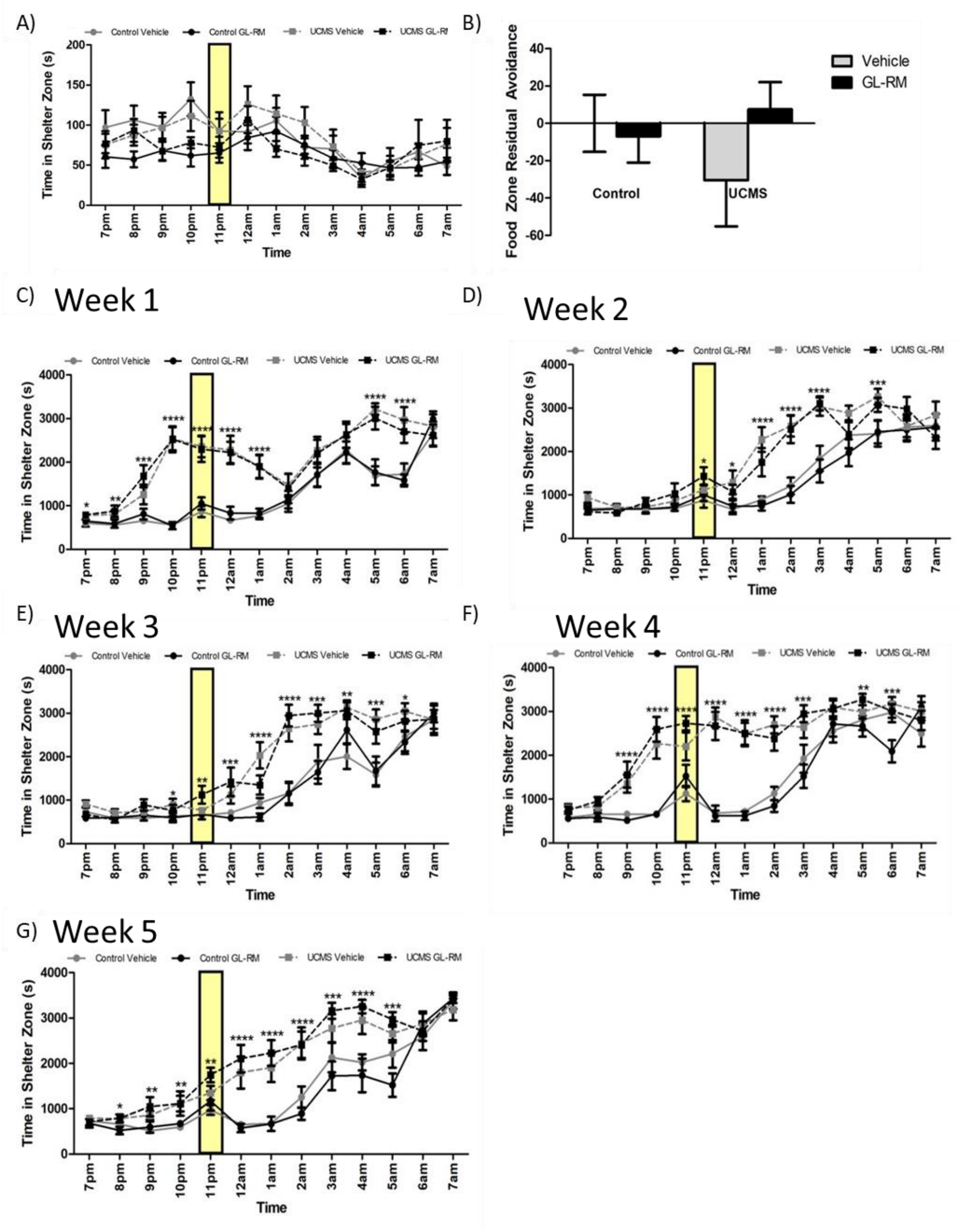

***Supplementary Figure 4.*** *Time spent in the shelter zone in the Phenotyper Boxes, at baseline and during UCMS.*

Prior to being exposed to UCMS and to treatment, mice were placed in the Phenotyper to measure their baseline activity, overnight from 7pm to 7am. A light challenge was applied from 11pm to 12am, above the food zone, creating an aversive stimulus. While animals were not yet subjected to chronic stress or injected with GL-RM, time spent in shelter zone (A) and respective residual avoidance (B), analyses did not identify any differences between groups. Then mice were placed in the Phenotyper on a weekly basis, during the course of the UCMS procedure (C-G). While animals were subjected to chronic stress, treatment with GL-RM was only applied on Week 6 (presented in Figure 2). Animals subjected to chronic stress spent more time in the shelter zone, compared to Control animals. *p<0.05, **p<0.01, ***p<0.001 compared to Control-Vehicle.

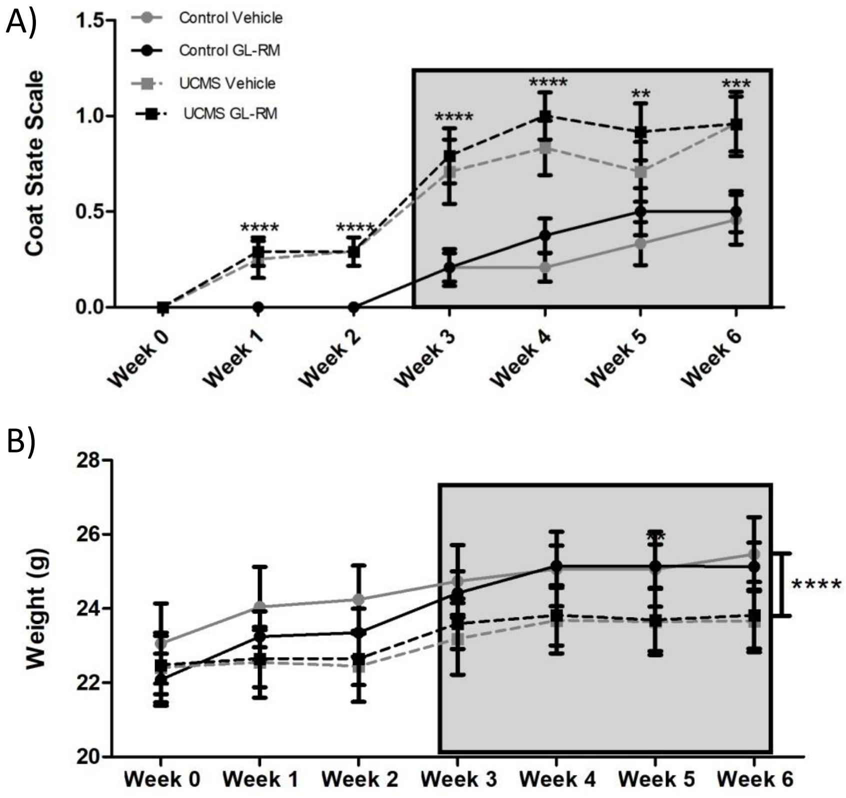

***Supplementary Figure 5.*** *Coat State and weight gain in mice from the chronic treatment study*

Coat state (A) and weight gain (B) were assessed on a weekly basis. Coat state showed a progressive increase in deterioration of the coat state in animals subjected to UCMS ($F_{(1;44)}$=23.848; p<0.0001), regardless of the treatment group ($F_{(1;44)}$=0.673; p=0.4164). Analyses of weight gain over the weeks showed an overall significant effect of stress ($F_{(1,44)}$=20.886; p<0.0001), and no effect of treatment ($F_{(1,44)}$=2.092; p=0.1552). Grey zone represents the period when animals were subjected to UCMS. **p<0.01, ***p<0.001 compared to Control-Vehicle.

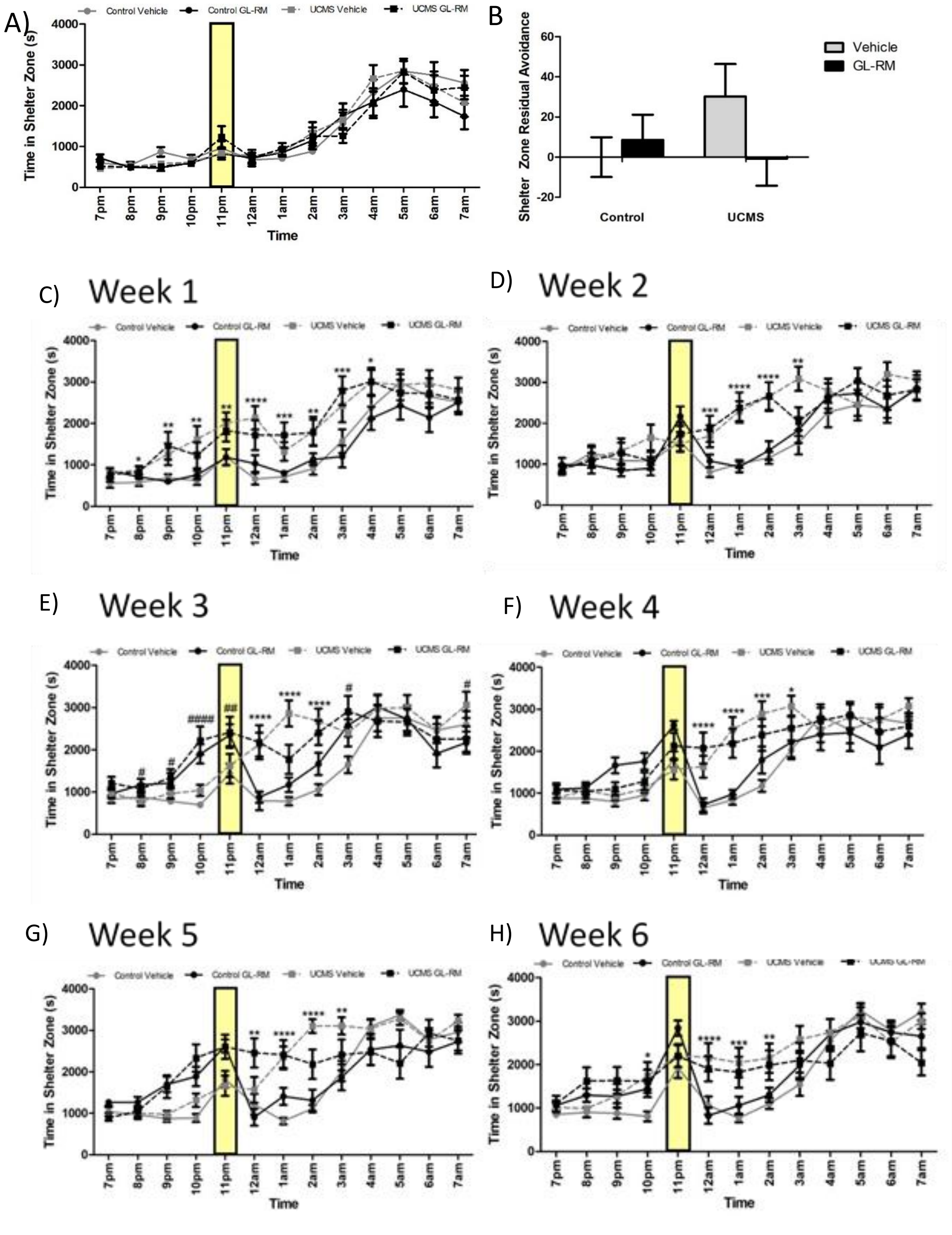
A)
Control Vehicle
Control GL-RM
UCMS Vehicle
UCMS GL-RM
Time in Shelter Zone (s)
Time
B
Shelter Zone Residual Avoidance
Vehicle
GL-RM
Control
UCMS
C)
Week 1
D)
Week 2
E)
Week 3
F)
Week 4
G)
Week 5
H)
Week 6

***Supplementary Figure 6.*** *Time spent in the shelter zone in the phenotyper, at baseline and during chronic stress and chronic treatment*

Mice were placed in the Phenotyper boxes prior to starting UCMS of treatment, to measure their activity overnight from 7pm to 7am. A light challenge was applied from 11pm to 12am above a food zone, creating an aversive stimulus. Time spent in the shelter zone (A) is presented here as well as Residual Avoidance (B). They were no differences between groups at baseline. Mice were then placed in the Phenotyper on a weekly basis. While animals were subjected to chronic stress, treatment with GL-RM was applied from Week 4 through Week 6. Animals subjected to chronic stress spent more time in the shelter zone, compared to Control animals, but there were no clear effect of GL-RM on overall time spent in the shelter Zone. *$p<0.05$, **$p<0.01$, ***$p<0.001$ compared to Control-Vehicle.

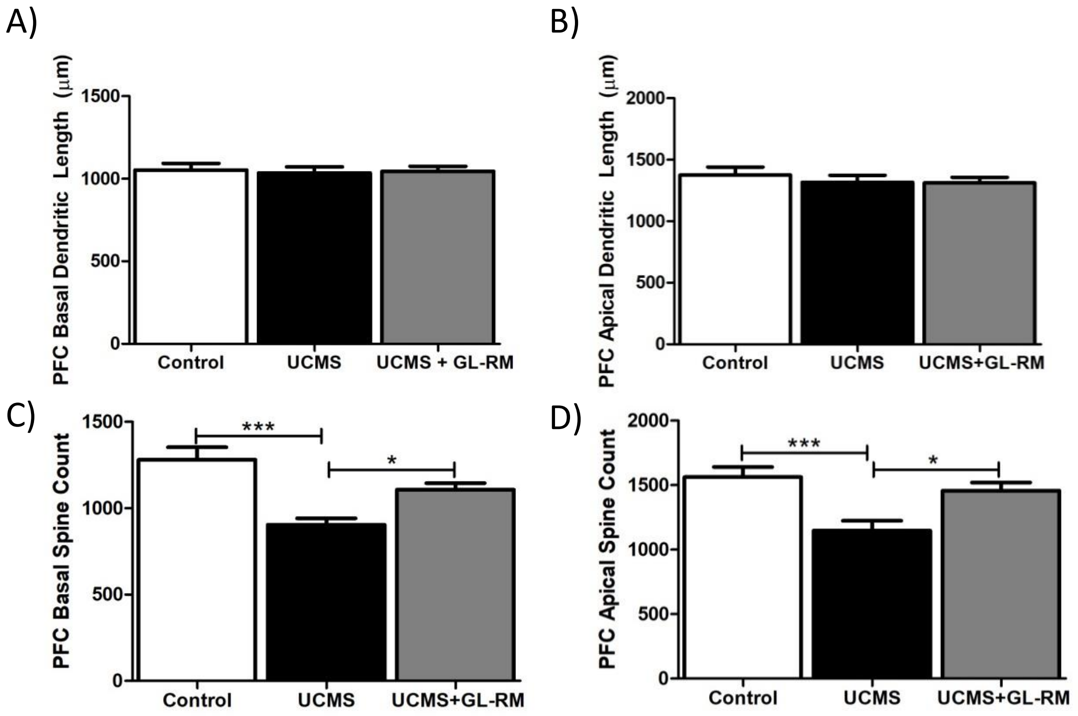

***Supplementary Figure 7.*** *Total dendritic length and spine count in the PFC of mice treatment with chronic GL-RM*

Brains were stained using a Golgi Cox technique and sectioned on a cryostat. Dendritic length and spine count in the prefrontal cortex were measured on 6 cells per animal, using a stereology-based approach with a Neurolucida software. No changes were observed in dendritic length in basal (A; $F_{(2;69)}$=0.06033; p=0.9415) nor apical (B; $F_{(2;69)}$=0.3887; p=0.6794) segments. ANOVA performed on spice count found group differences in both the basal and apical segments ($F_{(2;69)}$=25.28; p<0.0001, $F_{(2;69)}$=8.670; p=0.0004, respectively). This was due to asignificant decrease in basal (C) and apical (D) spine count in mice subjected to UCMS, that was reversed by chronic GL-RM treatment. *p<0.05, ***p<0.001 compared to “UCMS”.

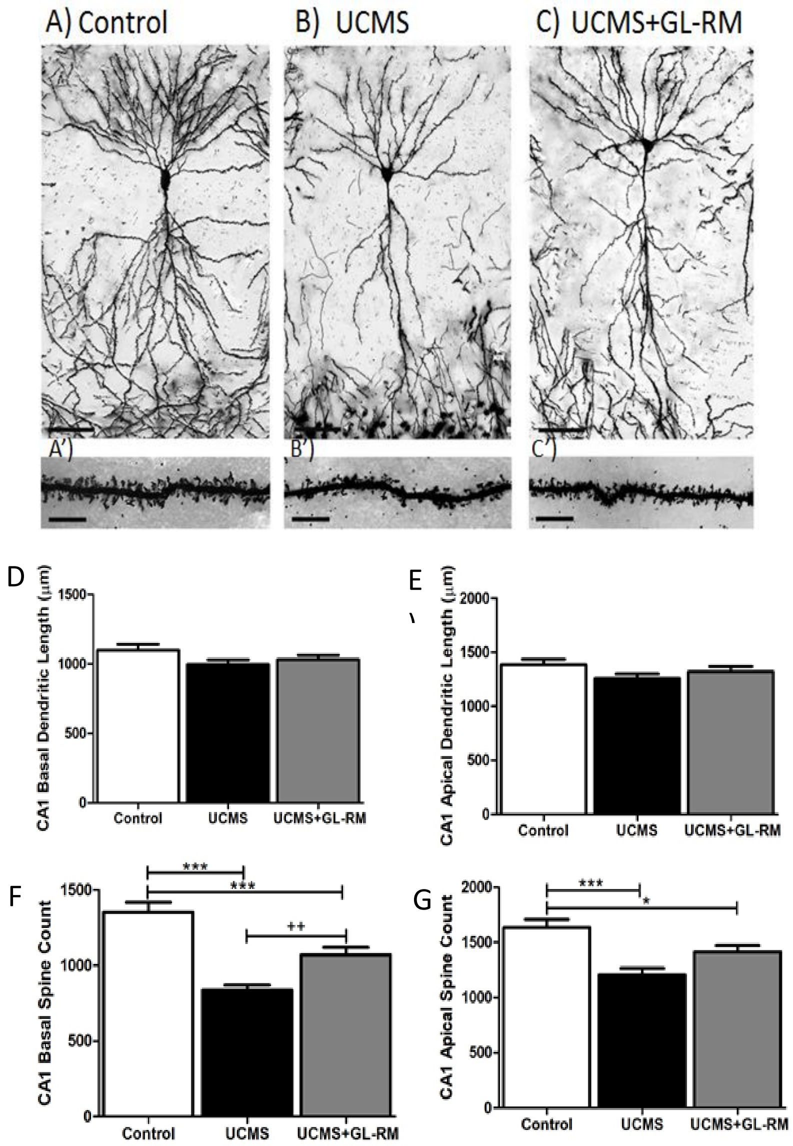

***Supplementary Figure 8.*** *Representative images of pyramidal neurons in the CA1 field of the hippocampus and total dendritic length and spine count in the CA1 of mice treatment with chronic GL-RM*

Brains from Control, UCMS and UCMS+GL-RM mice were harvested and stained using a Golgi-Cox technique. Hippocampal sections were mounted on slides and imaged at 100X for 3D dendritic reconstruction, followed by counting of the spines throughout the entire dendritic trees (A-C; Scale bar represents 50µm). Dendritic length and spine count in the CA1 of the hippocampus were measured on 6 cells per animal, using a stereology-based approach with a Neurolucida software. No changes were observed in dendritic length in the basal (D; $F_{(2;69)}$=2.202; p=0.1183) or apical (E; $F_{(2;69)}$=1.929; p=0.1530) segments. Significant differences between groups in spine count were identified in basal (F; ($F_{(2;69)}$=25.84;p=<0.0001) and apical (G; $F_{(2,69)}$=11.29;p<0.0001) segments. Mice subjected to UCMS had lower spine counts, that was partially reversed by chronic GL-RM treatment. *p<0.05, ***p<0.001 compared to “Control”, ++p<0.01 compared to “UCMS”
